# BLMM: Parallelised Computing for Big Linear Mixed Models

**DOI:** 10.1101/2022.03.09.483645

**Authors:** Thomas Maullin-Sapey, Thomas E. Nichols

## Abstract

Within neuroimaging large-scale, shared datasets are becoming increasingly commonplace, challenging existing tools both in terms of overall scale and complexity of the study designs. As sample sizes grow, researchers are presented with new opportunities to detect and account for grouping factors and covariance structure present in large experimental designs. In particular, standard linear model methods cannot account for the covariance and grouping structures present in large datasets, and the existing linear mixed models (LMM) tools are neither scalable nor exploit the computational speed-ups afforded by vectorisation of computations over voxels. Further, nearly all existing tools for imaging (fixed or mixed effect) do not account for variability in the patterns of missing data near cortical boundaries and the edge of the brain, and instead omit any voxels with any missing data. Yet in the large-*n* setting, such a voxel-wise deletion missing data strategy leads to severe shrinkage of the final analysis mask. To counter these issues, we describe the “Big” Linear Mixed Models (BLMM) toolbox, an efficient Python package for large-scale fMRI LMM analyses. BLMM is designed for use on high performance computing clusters and utilizes a Fisher Scoring procedure made possible by derivations for the LMM Fisher information matrix and score vectors derived in our previous work, Maullin-Sapey and Nichols (2021).

## 1. Introduction

### 1.1. Background

The field of functional Magnetic Resonance Imaging (fMRI) has recently seen a drastic improvement in terms of the volume of data collected and shared publicly. Many researchers now regularly face analyses involving “large-*n*” (large number of observations) datasets consisting of tens of thousands of images, typically endowed with some form of complex covariance structure (Smith and Nichols (2018), Li et al. (2019), Haworth et al. (2019)). For example, the Adolescent Brain Cognitive Development (ABCD) and UK Biobank (UKB) datasets, which contain imaging data from 10, 000 and 30, 000 subjects, respectively, both possess a multi-level covariance structure induced through a repeated measures experimental design (Casey et al. (2018), Allen et al. (2012)). Similarly, the well-known Enhancing NeuroImaging Genetics through Meta-Analysis (ENIGMA) cohort, which contains imaging data from tens of thousands of subjects, also possesses a multi-level covariance structure due to its pooling of data from many different sources (Bearden and Thompson (2017)). A more complex, but equally popular example is the Human Connectome Project (HCP) dataset, which contains observations drawn from 1200 subjects, but exhibits a constrained covariance structure due to relatedness between individuals and the sampling of family units (Van Essen et al. (2013)).

Often, covariance structures in an experimental design arise from grouping factors present in the data. Accounting for complex grouping factors during an fMRI analysis is an historically routine practice for small-sample studies. Commonly employed analysis designs in the small-sample setting include longitudinal multi-session analyses (observations grouped by subject), comparative group analyses (subjects grouped by study conditions) and mega-analyses (analysis results grouped according to study protocols). The widely-accepted, conventional approach to modelling such datasets is to employ the Linear Mixed Model (LMM) (c.f. Laird and Ware (1982), Friston et al. (2002)). The LMM accounts for complex grouping structures in datasets via the inclusion of both “fixed effects” and “random effects” during model specification. Fixed effects are unknown constant parameters that are associated with covariates in the experimental design. Random effects are random variables that model the systematic differences between instances of a categorical variable (e.g. between-subject differences, between-site differences).

There are many strong obstacles to the practical execution of LMM analyses for large-*n* fMRI datasets. Generally, an LMM analysis consists of two stages: parameter estimation and statistical inference, each of which presents unique computational and theoretical challenges in the large-*n* fMRI analysis setting. An overview of the challenges specific to each of these stages is provided in Sections 1.1.1 and 1.1.3, respectively. An additional issue, which merits separate consideration, concerning missing data found near cortical boundaries, is detailed in Section 1.1.2. In the univariate (non-imaging) setting, many tools exist for estimating the parameters of and performing inference upon the LMM (c.f. Section 2.1.3). However, many of these tools are not designed to (a) be scalable to arbitrarily large datasets and (b) exploit vectorisation speed-ups from processing multiple voxels simultaneously. To counter this, in recent work we derived novel closed form expressions for the Fisher information matrices and LMM gradient vectors (Maullin-Sapey and Nichols (2021)), making vectorised Fisher scoring practical for mass-univariate analysis. In this work, we shall demonstrate that these expressions may be employed to perform fast and scalable LMM parameter estimation and inference in the context of large-*n* fMRI analysis.

In this paper, we present “Big” Linear Mixed Models (BLMM), a Python-based tool for parameter estimation and statistical inference of mass-univariate LMMs. The BLMM tool partitions fMRI analyses to limit memory consumption, while still being able to exploit vectorization speed-ups from working with multiple voxels. Despite being built specifically for use on Sun Grid Engine (SGE) clusters, SGE-specific code in BLMM is isolated in one file and may be adapted for use on any HPC scheduler. In the following sections, we first provide background on LMM parameter estimation and inference as well as the “voxel-wise missingness” ubiquitous in large-*n* fMRI analyses. Following this, we give preliminary statistical information describing the univariate LMM and its extension to the mass-univariate voxel-wise setting. In the methods section, we then outline the computational pipeline of BLMM, starting with the input specification, followed by the distributed stages of computation, parameter estimation and finally, inference. Next, the correctness and performance of BLMM are assessed via simulation. We conclude by providing a real data example based on the UK Biobank.

#### 1.1.1. LMM Parameter Estimation

A vast amount of literature exists on the development of LMM parameter estimation tools and methodology. Primarily proposed in the late 1970s and early 1980s, many early approaches to LMM parameter estimation involved performing likelihood maximisation via numerical methods such as Fisher Scoring, Newton-Raphson and Expectation Maximisation (c.f. Dempster et al. (1977), Jennrich and Schluchter (1986), Laird et al. (1987)). More recently, several tools which build upon these fundamental ideas have become available for LMM parameter estimation in the univariate (single-model) setting. The most popular of these include the SAS and SPSS packages, PROC-MIXED and MIXED (SAS Institute Inc. (2015), IBM Corp (2015)), the R package lme4 (Bates et al. (2015)) and the Windows package HLM (S.W. et al. (2019)). These tools have been widely adopted within the statistical literature, largely due to their demonstrable computational efficiency when performing parameter estimation for a single univariate model.

However, in mass-univariate fMRI, parameter estimation is not performed for a single model, but rather for hundreds of thousands of models, each corresponding to a different voxel in the analysis mask. For a mass-univariate voxel-wise analysis to truly utilise the computational power available, it is a necessity that computation is vectorised across voxels. Many of the established LMM tools are reliant upon operations that are not naturally amenable to vectorisation. Examples of such operations include the sparse Cholesky decomposition employed by lme4 (Bates and DebRoy (2004)) and the sparse sweep operation employed by PROC-MIXED and MIXED (Wolfinger, Tobias, and Sall (1994)). The approaches employed by HLM circumvent this issue, but at the cost of generalizability since HLM only allows the estimation of LMMs which exhibit pre-specified structures, with an extension HCM providing additional options (Raudenbush (2002)). Operations that are not amenable to vectorisation create bottlenecks for mass-univariate computation as they must be executed separately for each voxel in the image. Serial execution of such operations can result in severe computational overheads, especially when modelling large sample sizes. As a result, many of the tools available for univariate LMM analysis cannot be employed in the large-*n* fMRI setting.

In the small-sample fMRI setting, several tools exist for mass-univariate LMM parameter estimation. These include SPM’s built-in mixed-effects module (Friston et al. (2005)), FSL’s FLAME package (Beckmann et al. (2003)), FreeSurfer’s longitudinal analysis pipeline (Bernal-Rusiel et al. (2013a)) and AFNI’s 3dLME package (Chen et al. (2013)). The tools provided by SPM, FSL and FreeSurfer perform parameter estimation for only LMMs in which observations are grouped by a single categorical factor. Specifically, SPM and FSL allow LMM parameter estimation of second-level designs (i.e. designs with subjects grouped by experimental features) whilst FreeSurfer allows for parameter estimation of longitudinal LMM designs (i.e. designs with timepoints grouped by subject) (FIL Methods Group (2020), Woolrich et al. (2004), Bernal-Rusiel et al. (2013b), Madhyastha et al. (2018)). In SPM and FreeSurfer, parameter estimates are obtained via Restricted Maximum Likelihood (REML) estimation, whilst in FSL a Bayesian approach is adopted. In contrast, AFNI’s 3dLME package allows parameter estimation of a much broader range of LMM designs and provides similar options to those offered by standard tools in the univariate setting. 3dLME provides this support by calling directly to the R package lme4 and parallelising computation across all available processors and cluster nodes via the use of the Dask Python package (Rocklin (2015)).

The tools provided by SPM, FSL, FreeSurfer and AFNI are computationally efficient for parameter estimation in the small-sample mass-univariate analysis setting, but were not originally designed for applications involving large sample sizes. In the context of large-*n* analyses, SPM, FSL and FreeSurfer quickly encounter memory errors as sample sizes increase into the hundreds whilst AFNI experiences overheads in terms of computation time. For all tools, reduced computational performance in the large-*n* setting primarily stems from two issues: (1) construction and storage of the analysis design (i.e. the design matrices and response vector) and (2) bottleneck computations which must be performed independently for each voxel.

#### 1.1.2. Missing Data

An important issue that must be addressed prior to or during the parameter estimation stage of an fMRI analysis is the missing data observed on and around cortical boundaries. Such missingness is ubiquitous in whole-brain fMRI analyses and can be attributed to several commonplace sources of between-image spatial variability. Such sources include magnetic susceptibility artefacts, imperfections in the image alignment process, differing image acquisition parameters and, indirectly, between-subject biological variation. Conventionally, standard fMRI analysis tools address this missing data problem by omitting voxels with incomplete observations from the analysis. As detailed by Vaden et al. (2012), and later by Gebregziabher et al. (2017), adopting this approach can negatively impact both the specificity and sensitivity of the analysis results, especially when spatial extent thresholding is employed for multiple comparisons correction. In terms of specificity, an inflated Type II error rate may be observed when the removal of voxels with incomplete data causes brain regions that are near cortical boundaries to be excluded from the analysis. In terms of sensitivity, an inflated Type I error rate may result from the smaller number of tests being performed, and consequently, the use of a less conservative multiple comparisons correction. Whilst the removal of missing-data voxels in the small-sample setting typically has a negligible effect, in the large-*n* setting omitting voxels can profoundly influence results of an analysis, often ‘deleting’ large chunks of the final images produced. The reason that the severity of this issue becomes notably more pronounced in the large-*n* setting is that the probability of a given voxel being missing in at least one image increases with the number of images in the analysis. To address this issue, the patterns of missingness observed for each voxel must be carefully considered and accounted for in large-*n* analyses.

#### 1.1.3. Inference

In the small-sample setting, fMRI LMM analyses conclude with significance-based hypothesis tests using Wald test statistics. However, hypothesis testing of this nature has long been a contentious topic in the broader LMM literature, due to unknown variability in the estimation of variance components and lack of exact distribution for the Wald test statistics (Verbeke and Molenberghs (2001)). As a result, whilst many of the popular tools for univariate LMM analysis provide support for calculating Wald T-statistics and F-statistics, there is debate concerning how the corresponding p-values are to be calculated and whether such practices should be widely adopted (c.f. Manor and Zucker (2004), Baayen et al. (2008), Luke (2017)).

The lack of consensus in the LMM literature is reflected by the range of options available for performing LMM inference in the univariate setting. For example, HLM, MIXED and PROC-MIXED each adopt the assumption that the Wald test statistics for the LMM ‘approximately’ follow student’s *t*- and *F*-distributions, but employ different techniques for estimating the associated unknown degrees of freedom. HLM approximates the degrees of freedom via closed-form expressions resembling those employed for multi-level linear model analyses (c.f. Raudenbush (2002), West et al. (2014)). On the other hand, MIXED and PROC-MIXED utilize the Welch-Satterthwaite equation (c.f. Section 2.1.4) and numerical gradient optimization (Satterthwaite (1946), Welch (1947), Fai and Cornelius (1996)) in order to obtain degrees of freedom estimates for a more general breadth of LMM applications. For brevity, in the remainder of this paper we shall refer to methods involving degrees of freedom estimation using the Welch-Satterthwaite equation as ‘WSDF’ (Welch-Satterthwaite Degrees of Freedom) based methods.

By employing this ‘approximate distribution’ assumption, HLM, MIXED and PROC-MIXED are able to provide support for significance-based hypothesis testing, outputting *p*-values alongside Wald statistics for a wide variety of LMMs. While lmer rejects any approximation and offers no p-values (Bates (2006)), the supporting lmerTest package (Kuznetsova et al. (2017)) provides p-values by using the same methods as MIXED and PROC-MIXED. Numerous studies have found that the WSDF-based approach employed by MIXED, PROC-MIXED and lmerTest is notably more robust to small sample sizes, unbalanced designs, covariance heterogeneity (when fitting the correct covariance structure) and non-normality than the approach adopted by HLM (c.f. Keselman et al. (1999), Schaalje et al. (2002), Kuznetsova et al. (2017), Luke (2017), Francq et al. (2019)). However, the computational burden of the numerical gradient estimation required for the WSDF-based approach is substantial and constitutes a significant obstacle to large-*n* LMM analysis in the mass-univariate fMRI setting.

Several tools are available in the small-sample fMRI setting for significance-based hypothesis testing via Wald statistics. As with LMM parameter estimation, these include SPM’s built-in mixed-effects module, FSL’s FLAME package, FreeSurfer’s longitudinal analysis pipeline and AFNI’s 3dLME package. Due to the low computational costs required, SPM, FreeSurfer and FSL each employ a similar approach to that used by HLM in the univariate setting by using closed-form expressions, which can be found, for example, in Pinheiro and Bates (2009), to approximate the degrees of freedom. AFNI alternately employs the WSDF approach by acting as a wrapper for the lmerTest package. While the former approach is more efficient in terms of computation time and memory, it provides less accurate estimates for the degrees of freedom. Conversely, the latter approach provides more accurate estimation of the approximate sampling distributions of the Wald test statistics, but at the cost of increased computation time that scales with the number of observations. In this work, we make use of recent novel closed-form expressions we developed for evaluating the gradients required by the WSDF approach (Maullin-Sapey and Nichols (2021)). These expressions offer a viable and accurate alternative to gradient estimation and may be employed in the large-*n* fMRI setting using vectorised computation without sacrificing statistical accuracy.

### 1.2. Preliminaries

In this section, a brief overview and statement of the mass-univariate LMM is provided. To simplify notation, we begin by defining the univariate LMM in Section 1.2.1. Following this, in Section 1.2.2 we describe how the definition and notation of Section 1.2.1 are extended to the mass-univariate fMRI setting.

#### 1.2.1. The Linear Mixed Model

In the traditional univariate setting, an LMM containing *n* observations is assumed to take the following form:

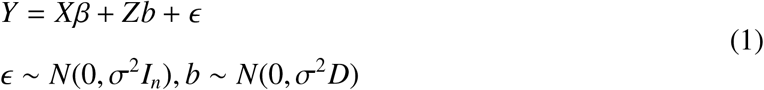

where the observed quantities are the response vector *Y*, fixed effects design matrix *X*, and random effects design matrix *Z*, and the unknown model parameters are the fixed effects parameter vector β, the scalar fixed effects variance *σ*^2^, and the random effects covariance matrix *D*.

The random effects in the model are specified using factors (categorical variables which group the random effects) and levels (the individual instances of the categorical factors). The total number of factors that group the random effects in the model is denoted as *r*. For the *k*^*th*^ factor in the model, *l*_*k*_ and *q*_*k*_ are used to denote the number of levels belonging to the factor and the number of random effects that the factor groups, respectively. The random effects design matrix, *Z*, can be partitioned horizontally as 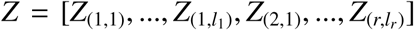 where *Z*_(*k, j*)_ consists of the random effects covariates which are grouped into the *j*^*th*^ level of the *k*^*th*^ factor in the model. The random effects covariance matrix, *D*, is block diagonal and can be specified as 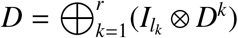, where ⊕ is the direct sum and *D*^*k*^ is the (*q*_*k*_ × *q*_*k*_) within-level covariance matrix for the *k*^*th*^ factor in the model. This notation is essential for describing the Fisher Scoring algorithm approach that will be employed by BLMM for parameter estimation (see Section 2.1.3).

From equation (1), the restricted log-likelihood function for the LMM, ignoring constant terms, is given by:

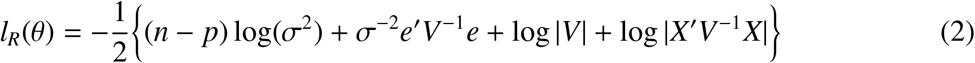

where *θ* is shorthand for the parameters (*β, σ*^2^, *D*), *p* is the number of fixed effect parameters in the design, *V* = *I*_*n*_ + *ZDZ*′ is the marginal variance, and *e* = *Y* − *Xβ* is the residual vector. Throughout this work, we assume that *θ* takes the form *θ* = [*β*′, *σ*^2^, vec(*D*^1^)′, …, vec(*D*^*r*^)′]′, where vec represents the vectorization operator. It should be noted that whilst this may seem like a natural representation of *θ*, it is by no means the only possible representation. A full discussion of this, alongside a more detailed introduction to the LMM and the notation described in this section, is provided in our previous work, Maullin-Sapey and Nichols (2021).

#### 1.2.2. The Mass Univariate Model

In the mass-univariate setting we fit and infer on hundreds of thousands of LMMs concurrently. In the setting of fMRI, each LMM corresponds to a voxel in the study’s analysis mask. Adapting the notation of the previous section, this is represented as:

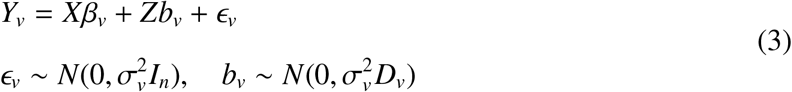

where the subscript *v* represents voxel number. In equation (3), the fixed effects and random effects design matrices (*X* and *Z*) are treated as constant across all voxels, whilst the response vector (*Y*), design parameters (*β, σ*^2^ and *D*) and random terms (*ϵ* and *b*) vary from voxel to voxel. By extension, this also means that *n, r*, {*l*_*k*_}_*k*∈{1,…*r*}_ and {*q*_*k*_}_*k*∈{1,…*r*}_ are also treated as constant across voxels. Equation (3) is the conventional form of the mass-univariate LMM which is typically adopted by the existing fMRI LMM software packages. As noted in Section 1.1.2, this model does not account for mask-variability and, as a result, must be adapted to reflect the pattern of missingness observed at each voxel.

To account for such missingness in *Y*_*v*_, we adopt an MCAR (Missing Completely At Random) assumption and define the (*n* × *n*)-dimensional ‘missingness matrix’, *M*_*v*_, as a diagonal indicator matrix, where the (*i, i*)^*th*^ element is 1 if the *i*^*th*^ element of *Y*_*v*_ is not missing and 0 otherwise. We now define *X*_*v*_ = *M*_*v*_*X* and *Z*_*v*_ = *M*_*v*_*Z* and assume that missingness in *Y*_*v*_ and *ϵ*_*v*_ is encoded as 0. The model specification for a mass-univariate LMM that accounts for missingness induced by mask-variability is now given as follows:

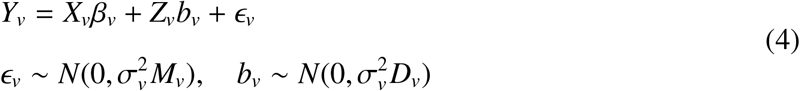

The inclusion of the missingness matrix, *M*_*v*_, ensures that rows of the design at which missingness occurred for voxel *v* are replaced with zeros, or “zero-ed out”. This process ensures that the analysis proceeds as though such ‘missing-data’ rows had not been included in the design at all.

An important ramification of this model construction is that the fixed effects and random effects design, *X*_*v*_ and *Z*_*v*_, as well as the number of observations, *n*_*v*_, are now treated as spatially varying. If naively implemented, a model involving a spatially varying design (such as equation (4)) presents a much more formidable computational challenge than its non-spatially varying counterpart (i.e. equation (3)), as additional expenses in memory and computation time arise from accounting for voxel-specific design matrices. However, in this context, it must be noted that as *X*_*v*_ and *Z*_*v*_ are constructed by ‘zero-ing out’ rows of *X* and *Z*, many voxels will employ identical design matrices for analysis (i.e. it is often the case that 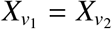 and 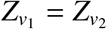 for two separate voxels *v*_1_ and *v*_2_). In particular, it is expected that inside the brain, far from cortical boundaries, no missingness will be observed, and therefore, for many voxels, *X*_*v*_ = *X* and *Z*_*v*_ = *Z*. In other words, whilst it is true that the design matrices, *X*_*v*_ and *Z*_*v*_, vary spatially across the whole brain, it is expected that large contiguous groups of voxels will exist over which *X*_*v*_ and *Z*_*v*_ do not vary at all. Accounting for this “between-voxel design commonality” can result in drastic improvements in computation speed and efficiency, and thus, constitutes a key motivation behind the approach adopted by BLMM.

In the following sections, unless stated otherwise the subscript *v*, representing voxel number, will be dropped from our notation. For the remainder of this work, it is assumed implicitly that any equations provided correspond to a model of the form (4) for some given voxel.

## 2. Methods

In this section, we describe the computational pipeline employed by BLMM to perform mass-univariate LMM analysis of large-*n* fMRI data, as well as the simulations and real-data examples for which results are later presented in Section 3. To begin, Section 2.1 gives an in-depth overview of the stages of the BLMM computational pipeline. Following this, Section 2.2 describes simulations that shall be used to assess the correctness and performance of BLMM. Finally, Section 2.3 details a real-world example based on repeated-measures data drawn from the UK Biobank, demonstrating BLMM’s usage in practice.

### 2.1. The BLMM Pipeline

A visual overview of the BLMM pipeline is provided by Fig. 1 in the form of an activity diagram. Highlighted in Fig. 1 are the four “stages” of the BLMM pipeline: input specification, product form computation, parameter estimation, and inference and output. Each of these stages is described in turn by Sections 2.1.1-2.1.4, respectively. Also labelled in Fig. 1 are the steps of the pipeline which employ distributed computation: “Image-wise batching” and “Voxel-wise batching”, each of which is further discussed further in Sections 2.1.2 and 2.1.3-2.1.4, respectively.

**Figure 1:**
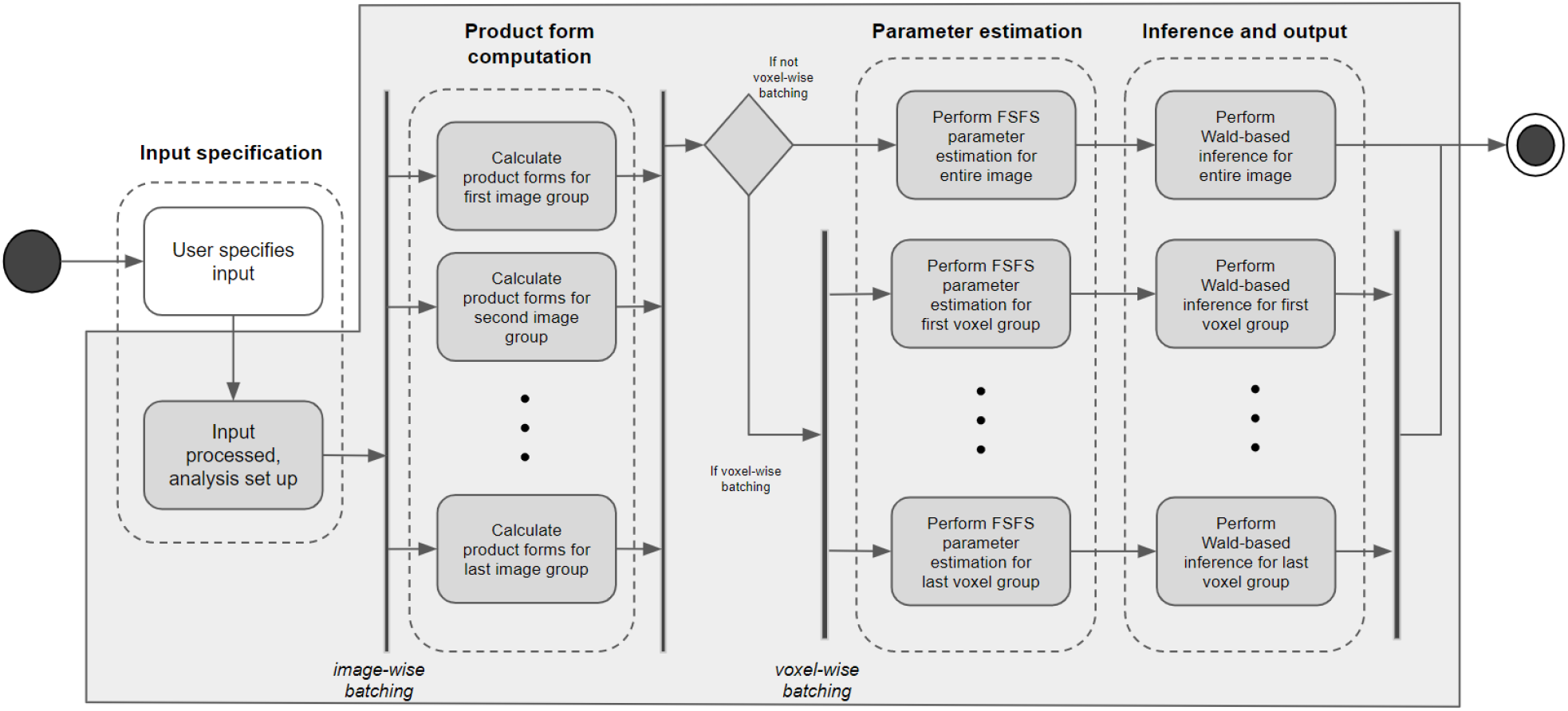
Activity diagram detailing the BLMM pipeline. The boundary of the BLMM code is indicated by the gray outline. The start and end nodes of the pipeline are represented by the black circle and nested black and white circles, respectively. Decision nodes are represented by diamonds and parallel stages of computation are represented with vertical bars. Also included are dotted lines indicating the distinct “stages” of the BLMM pipeline.

#### 2.1.1. Input Specification

To specify an analysis design in BLMM, the user must provide the fixed effects design matrix, *X*, the response images, *Y*, and the random effects design matrix, *Z*. For specification of the random effects design matrix, *Z*, a similar approach to that of lmer is adopted (Bates et al. (2015)). For each factor in the design, the user provides a factor vector, *g*_*k*_, and a “raw” regressor matrix, *z*_*k*_. The raw regressor matrix, *z*_*k*_, is a matrix of dimension (*n* × *q*_*k*_) and contains the covariates which correspond to the random effects that are to be grouped by the *k*^*th*^ factor in the model. The factor vector, *g*_*k*_, is a numerical vector of dimension (*n* × 1) with entries indicating to which level of the *k*^*th*^ factor each observation belongs. From these inputs, the following construction is used to obtain the random effects design matrix *Z*:

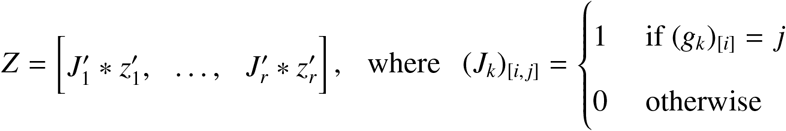

and * is the Khatri-Rao product. For example, in a longitudinal design, *g*_1_ would be a vector of subject identifiers and *z*_1_ could contain a column of 1’s for a random intercept and a column of study times for a random slope for the time effect. See Supplementary Material Section S1 for a complete worked example.

During input specification, BLMM also provides a range of masking options. By default, the user is required to specify an analysis mask. In addition, the user may specify one mask per input image, as well as a “missingness threshold”. The missingness threshold, which may be specified as either a percentage or an integer, indicates how many input images a voxel must have recorded data for in order for it to be retained in the final analysis. This threshold is essential, as while accommodating missingness is an essential feature of BLMM, allowing excessive missingness (down to a small faction of the data) is not advised and may result in rank-deficient designs. Implicit masking is also supported by BLMM, with any voxel set to 0 or NaN in an input image being treated as ‘missing’ from the analysis.

Whilst specifying the analysis model, the user may also opt to perform hypothesis testing using an approximate Wald *T* −test or *F*−test. To specify a hypothesis test of this form, the user must provide a contrast vector, *L*, representing the null hypothesis, and the type of statistic to be used to perform the test (i.e. *T* or *F*). Further detail on hypothesis testing via approximate Wald statistics is provided in Section 2.1.4.

#### 2.1.2. Product Form Computation

Computation within the BLMM pipeline begins by, for each voxel *v*, computing the “product forms” defined as follows:

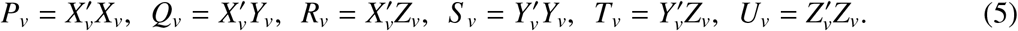

Following the computation of the product forms, the original matrices, *X*_*v*_, *Y*_*v*_ and *Z*_*v*_, can be discarded since only the product forms are required for future computation (c.f. Sections 2.1.3-2.1.4). This approach is adopted by BLMM as the dimensions of the product forms do not scale with *n* but rather with *p* and *q*, the second dimensions of the fixed effects and random effects design matrices, respectively. As *p* and *q* are assumed to be much smaller than *n*, working with the product forms instead of *X*_*v*_, *Y*_*v*_ and *Z*_*v*_ can provide large reductions in both memory consumption and computation time (c.f. Maullin-Sapey and Nichols (2021)).

To compute the product forms efficiently, BLMM employs an image-wise batching approach. In this approach, the input images and model matrices are split into batches and computation is performed in parallel across several cluster nodes. More precisely, given *B* nodes, *X* and *Z* are vertically partitioned into evenly sized blocks {*X*^(*b*)^}_*b*∈{1,…,*B*}_ and {*Z*^(*b*)^}_*b*∈{1,…,*B*}_, respectively. The list of input images, *Y*, is also partitioned into *B* lists of 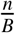 input images, {*Y*^(*b*)^}_*b*∈{1,…,*B*}_, each corresponding to a partition of *X* and *Z*. Each node is assigned a partition of *X*, the corresponding partition of *Z* and the corresponding list of input images. For every voxel in the analysis mask, this means that the *b*^*th*^ node now possesses 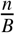 observations (assuming missing values are encoded as zero). We denote the response vector of observations at voxel *v*, taken from the images *Y*^(*b*)^, as 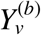.

The *b*^*th*^ node now applies voxel-specific masking to *X*^(*b*)^ and *Z*^(*b*)^ to obtain the spatially-varying designs 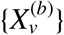 and 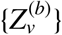. For each voxel, *v*, this is achieved by considering whether *v* has missing data in the input images listed in *Y*^(*b*)^ and “zero-ing” out the corresponding rows of *X*^(*b*)^ and *Z*^(*b*)^ accordingly (c.f. Section 1.2.2). Given the spatially varying designs, 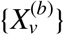 and 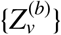, and response vector, 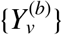, the product forms for this partition of the model are now computed as: 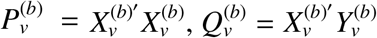,… and so forth.

For each voxel, the product forms for each partition are then sent to a designated central node. For the *v*^*th*^ voxel, the central node now computes the product forms for the entire model by summing over those sent from each node (i.e. 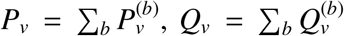,…). This approach can be seen to produce the product forms which were defined by equation (5) by noting, for arbitrary matrices of appropriate dimension, *A* and *B*, with corresponding vertical partitions {*A*^(*b*)^} and {*B*^(*b*)^}, that 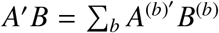. Pseudocode for the product form computation stage of the BLMM pipeline is provided by Algorithm 1.

To prevent convergence failure due to rank deficiency, following product form calculation, any voxels for which rank(*P*_*v*_) < *p* or rank(*U*_*v*_) < *q* are dropped from the analysis. Removal of such voxels is advised during analysis but can be prevented by setting the “safeMode” option to 0. The approach described in this section was initially motivated by a similar method employed for parameter estimation of the Linear Model. Details of this method are provided in Supplementary Material Section S2, alongside a corresponding implementation written in Python. Further notes on the computational efficiency of Algorithm 1 are also provided in Supplementary Material Section S3.

#### 2.1.3. Parameter Estimation

The most computationally intensive stage of any LMM analysis is the estimation of the unknown model parameters (*β, σ*^2^, *D*). A common approach to estimating the unknown model parameters of the LMM is to perform Restricted Maximum Likelihood (REML) estimation using Equation (2). BLMM employs the Full Simplified Fisher Scoring (FSFS) algorithm to perform this task for each voxel in the analysis mask. Proposed for the multi-factor LMM in our previous work, the FSFS algorithm iteratively performs updates to the fixed effects parameter vector, *β*, and fixed effects variance estimate, *σ*^2^, using the Generalized Least Squares (GLS) estimators, and to {vec(*D*^*k*^)}_*k*∈{1,…,*r*}_ separately using a Fisher Scoring update step based on the “full” representation of *D*^*k*^, vec(*D*^*k*^). (“Full” refers to the parameterization of vec(*D*^*k*^); see Section 2.1.2 of Maullin-Sapey and Nichols (2021) for further detail). Formally, during each iteration, *β* and *σ*^2^ are updated according to the following GLS update rules:

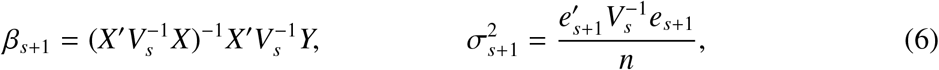

where *V*_*s*_ = *I* + *ZD*_*s*_*Z′, e*_*s*_ = *Y* − *Xβ*_*s*_ and the subscript *s* here, and throughout the remainder of this section only, denotes iteration number. For *k* ∈ {1, …, *r*}, the update rule employed for *D*^*k*^ takes the following form:

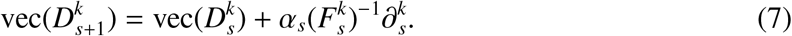

##### Algorithm 1: Product Form Computation Pseudocode

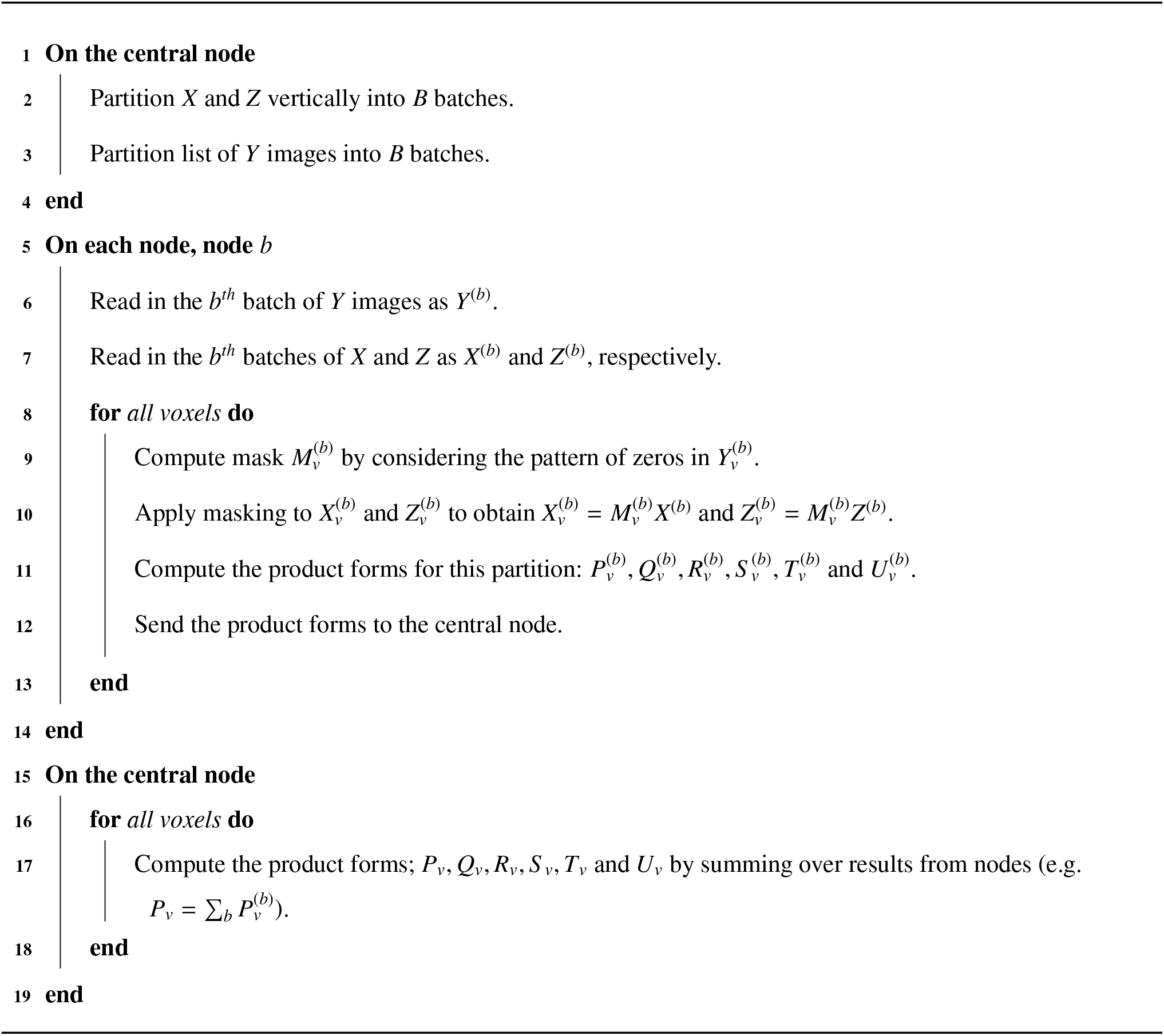

Here, *α*_*s*_ is a scalar step size, which is initialized to *α*_0_ = 1 and halved each time a decrease in log-likelihood is observed between iterations, 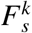 acts as a Fisher Information matrix given by:

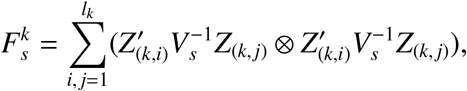

and 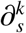 is the score vector given by:

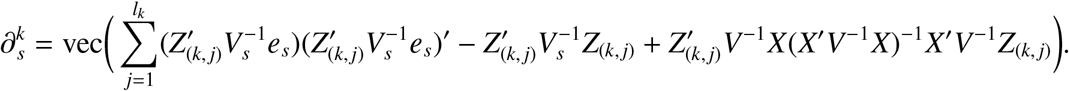

In this approach, to ensure that the estimates of {*D*^*k*^}_*k*∈{1,…*r*}_ are non-negative definite following each evaluation of the above update rules, an eigendecomposition based approach is used to project *D*^*k*^ to the space of non-negative definite matrices. Further detail on the use of the eigendecomposition in this manner can be found, for example, in Demidenko (2013). We note here that *F*^*k*^ is technically not a Fisher Information matrix, but rather a simplified version of the true Fisher Information matrix for the “full” representation of *θ* that provides the exact same updates during optimisation. For further details on this distinction, see our previous work, Maullin-Sapey and Nichols (2021).

It is important to note that, in every equation given above, the right-hand side can be reformulated to be expressed solely in terms of the product forms and the parameter estimates (*β, σ*^2^, *D*). This is crucial to the BLMM framework as, as is noted in Section 2.1.2, to prevent memory consumption and computation time from scaling with *n*, only the product forms are retained in memory following product form computation. The GLS update rules for *β* and *σ*^2^, equation (6), can be written in terms of the product forms and parameter estimates as follows:

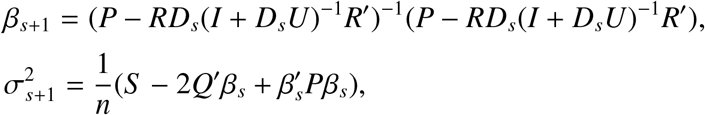

and the expressions for 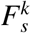 and 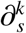 can be seen to be composed entirely of sub-matrices of 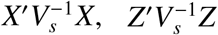 and 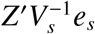, which are given by:

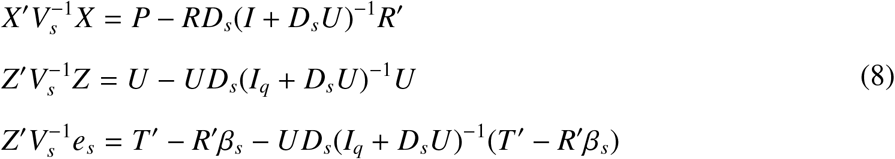

The restricted log-likelihood function, given by equation (2), can similarly be rewritten in terms of product forms as follows:

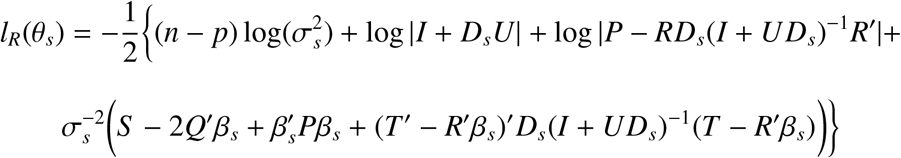

To choose initial values for the optimization procedure, BLMM follows the recommendations of Demidenko (2013) and Maullin-Sapey and Nichols (2021), employing the OLS estimators as starting estimates for *β* and *σ*^2^;

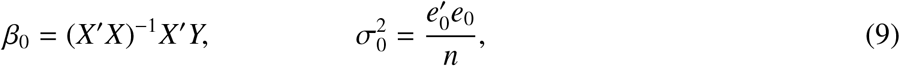

and the FSFS update rule, (7), with *I*_*n*_ substituted in the place of *V*, for a starting estimate of vec(*D*^*k*^):

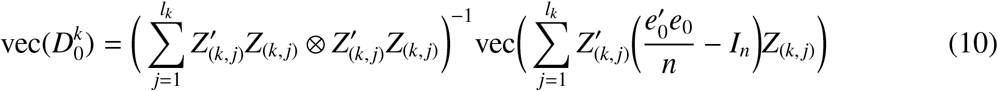

Again, the above expressions may be evaluated using only the product forms and do not require use of the original matrices *X, Y* and *Z*. Successful convergence of the FSFS algorithm occurs when the difference in log-likelihood observed between successive iterations becomes less than a predefined tolerance (10^−6^ by default) whilst convergence failure is deemed to have occured if the maximum number of iterations (10^4^ by default) is exceeded. The predefined tolerance and maximum number of iterations may be changed by the user via the “tol” and “maxnit” options, respectively. We note here that the default value of 10^4^ for maxnit is likely over-cautious as the simulations in our previous work (Maullin-Sapey and Nichols (2021)) found that, for a range of well-specified designs, the FSFS algorithm typically converged within 5 − 30 iterations.

Pseudocode for the FSFS algorithm is provided by Algorithm 2. In certain instances, further improvements in terms of computational performance can be obtained by utilising structural features of the analysis design which simplify the above expressions. In particular, BLMM has been optimized to give faster performance for models that contain (i) one random effect grouped by one random factor and (ii) multiple random effects grouped by one random factor. Further detail and discussion of the improvements employed for such models can be found in Supplementary Material Section S4.

##### Algorithm 2: Full Simplified Fisher Scoring Pseudocode

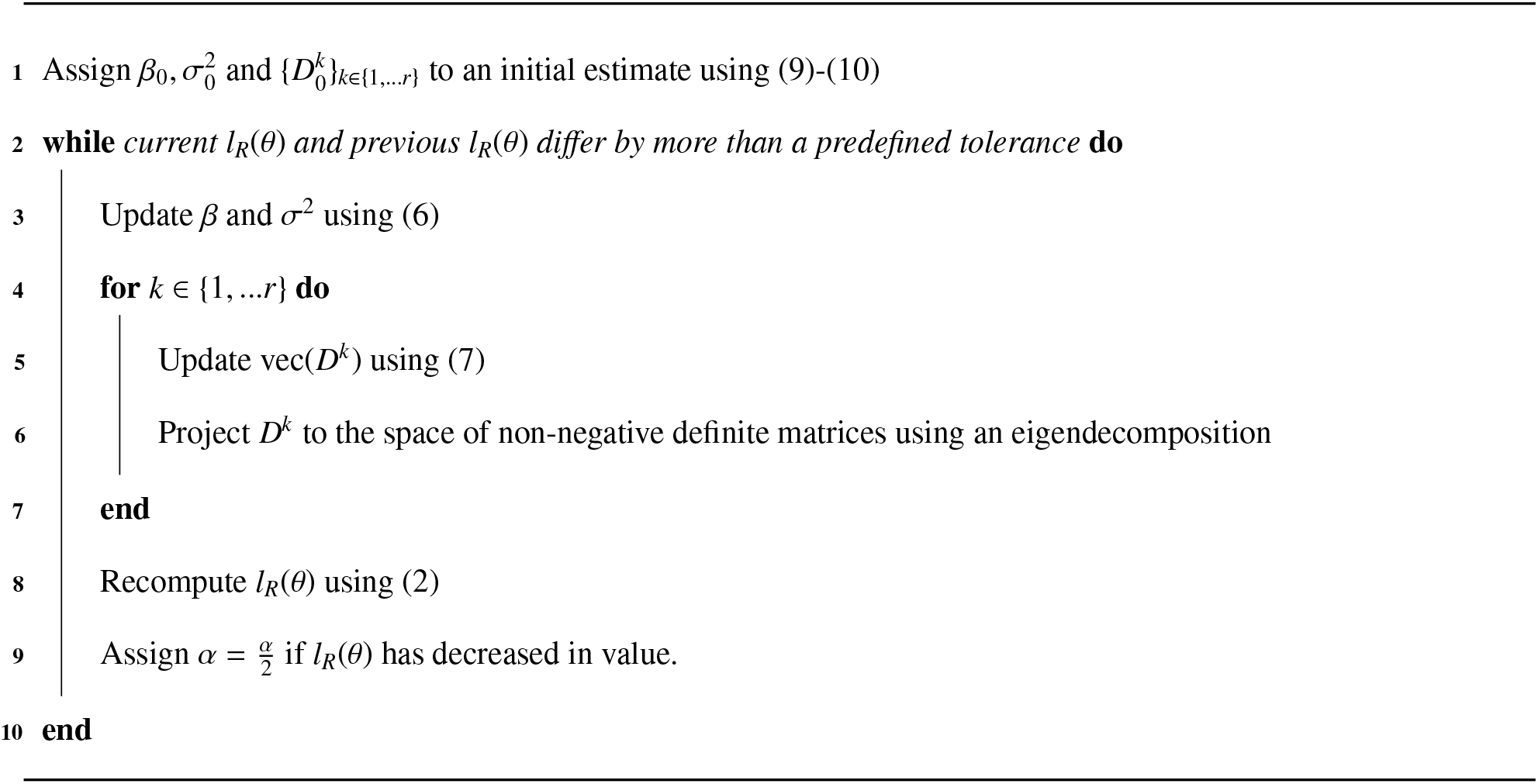

In the fMRI analysis setting, careful attention must be given to computational efficiency during parameter estimation, as parameters must be estimated for every voxel in the analysis mask. As noted in Section 1.1.1, computation that is performed independently for one voxel at a time can result in prohibitively slow computation speeds. It follows that, in order to execute LMM parameter estimation for fMRI data in a practical time frame, computation must be streamlined to allow for parameter estimation to be performed concurrently across multiple voxels at once.

On a single node, parameter estimation may be parallelised across voxels by using broadcasted computation, which exploits the repetitive nature of simplistic operations to streamline calculation. A considerable advantage in using the FSFS algorithm is that it relies upon only conceptually simplistic operations (such as matrix multiplication, matrix inversion and the eigendecomposition), for which a wealth of broadcasted support already exists in modern programming languages such as MATLAB and Python. By utilizing this support, the FSFS algorithm may be executed for multiple voxels concurrently in order to achieve quick and efficient computational performance (c.f. Section 3.1 for an assessment of BLMM computation time).

If multiple nodes are available, parameter estimation may also be parallelised further by partitioning the analysis mask into “batches” of voxels and having each node perform parameter estimation for an individual batch. This approach is also provided by BLMM and is referred to as “voxel-wise batching”. The ability to distribute computation in this manner is advantageous in situations where the analysis design is large and the product forms cannot be read into memory for many voxels at once. However, this additional layer of parallelisation may not be necessary for smaller designs and is therefore offered optionally in the BLMM package (as shown in Fig. 1).

#### 2.1.4. Inference and Output

The final stage of the BLMM pipeline is to perform inference on the fixed effects parameters and output the analysis results in NIfTI format. To support null-hypothesis testing for the fixed effects parameter vector, *β*, BLMM adopts an approach that is similar to that taken by the popular univariate LMM packages lmerTest, MIXED and PROC-MIXED (c.f. Section 1.1.3). In this approach, the REML estimates of the parameter vector, 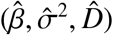, are used to construct Wald test statistics. To obtain corresponding *p*-values, a WSDF-based approach is then employed to model the sampling distribution of the Wald statistics.

Assuming the user has provided a contrast vector, *L*, to specify a null hypothesis *H*_0_ : *Lβ* = 0, BLMM will compute the corresponding Wald T-statistic or F-statistic for each voxel as follows:

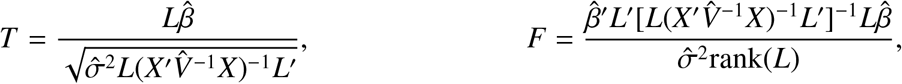

where 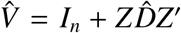. BLMM assumes that the distributions of *T* and *F* are reasonably approximated with a student’s *t*- or *F*-distribution. As the distributional assumptions for *T* and *F* are approximate, and not exact, the degrees of freedom are unknown and must be estimated. The estimation is performed using a WSDF-based approach based on the Welch-Satterthwaite equation;

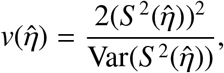

Where 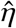 represents an estimate of the variance parameters η = (*σ*^2^, *D*^1^, … *D*^*r*^) and 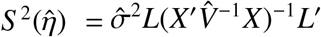. The numerator of the above expression may be evaluated directly. However, the below approximation, obtained using a second order Taylor expansion, must be employed to estimate the unknown variance term in the denominator:

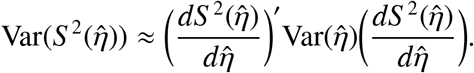

While other tools (lmerTest, MIXED, PROC-MIXED) require numerical optimisation to obtain this denominator, we make use of our previous work, Maullin-Sapey and Nichols (2021), where we presented novel closed-form expressions which may be used to evaluate the variance and derivative terms in the above directly. These expressions not only provide a computationally efficient alternative to the use of numerical optimisation, but also can be expressed purely in terms of the product forms (see Supplementary Material Section S5 for further detail). As a consequence of this, using a similar approach to that of Section 2.1.3, BLMM is able to perform degrees of freedom estimation by utilising only the product forms and parameter estimates at each voxel, and by employing operations that can be broadcasted or further accelerated through batch-wise parallelisation.

Once an analysis has been executed using BLMM, parameter estimate images for *β, σ*^2^ and *D*, contrast images of the form 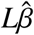 and Wald statistic images with corresponding *p*−value images are output, which must then be corrected for multiple testings. Correction for multiple testing is essential, though the non-linear estimation process precludes the use of standard random field theory and, while permutation or wild bootstrap methods are available for mixed models, such methods add substantial computational burden. Hence, either control of the Family-Wise Error rate (FWE) with the Bonferroni correction (Dunn (1961)) or the False Discovery Rate (FDR) with the Benjamini-Hochberg method (Benjamini and Hochberg (1995)) should be used to account for multiple testing.

Following the conclusion of a BLMM analysis, BLMM also provides Likelihood Ratio Testing (LRT) for comparison of LMM analyses which contain a single-random factor and differ only by the inclusion of random effects. More precisely, the output from several BLMM (or BLM, c.f. Supplementary Material Section S2) analyses may be used to perform hypothesis testing of the below form:

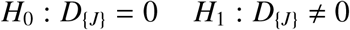

where *J* represents a predetermined set of elements of *D* which correspond to the removal of 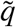 random effects from the study design. The above hypothesis is tested using the LRT statistic of the form 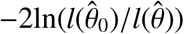, where 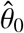 and 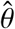 are the parameter estimates obtained from BLMM analyses in which the random effects *D*_{*J*}_ are and are not included in the model specification, respectively. Following the recommendations of Verbeke and Molenberghs (2001), BLMM assumes that the LRT test statistic follows the distribution 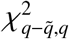, where 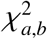 represents an even mixture of the distributions 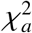 and 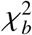. Under this distributional assumption, BLMM may be used to generate uncorrected p-value significance images for the LRT statistic. Examples of this method in practice are provided by Section 3.2.

### 2.2. Simulation Methods

In order to quantitatively assess and demonstrate the computational accuracy and efficiency of BLMM, extensive simulations were conducted. Simulated data was generated for nine simulation settings: three sample sizes (*n* = 200, 500 and 1000, respectively), each generated under three experimental designs. The experimental designs considered for simulation in this work reflect the two settings for which parameter estimation in BLMM has been explicitly optimized (c.f. Section 2.1.3 and Supplementary Material Section S4), as well as the most general model specification that BLMM caters to. These were the settings in which the experimental design contains; (i) one random factor which groups one random effect, (ii) one random factor which groups multiple random effects and (iii) multiple random factors, respectively. These models correspond to common use cases designs employing, for example, (i) a subject-level random intercept in a repeated measures setting, (ii) a subject-level random intercept and slope in a repeated measures setting, and (iii) a subject-level intercept and site-level intercept for repeated measures taken from multiple subjects across multiple sites. For each simulation setting, 1000 individual simulation instances were performed, and all reported results are given as averages taken across the 1000 instances.

In each simulation setting, the fixed effects parameter vector, *β*, and the fixed effects variance, *σ*^2^, were fixed across simulation instances and given by *β* = [4, 3, 2, 1, 0]′ and *σ*^2^ = 1, respectively. The fixed effects design matrix, *X*, contained an intercept and four regressors, each of which varied across simulation instances and consisted of values generated according to a uniform [−0.5, 0.5] distribution. Each experimental design enforced a different structure on the random effects design and covariance matrices, *Z* and *D*. The first experimental design (Design 1) included a single factor which grouped one random effect into 100 levels (i.e. *r* = 1, *q*_1_ = 1 and *l*_1_ = 100). The second experimental design (Design 2) included a single factor which grouped two random effects into 50 levels (*r* = 1, *q*_1_ = 2, *l*_1_ = 50). The third experimental design (Design 3) included two crossed factors, the first of which grouped two random effects into 20 levels and the second of which grouped one random effect into 10 levels (i.e. *r* = 2, *q*_1_ = 2, *q*_2_ = 1, *l*_1_ = 20 and *l*_2_ = 10). In all simulation instances, the first random effect appearing in *Z* was an intercept. Any additional random effects regressors varied across simulation instances and were generated according to a uniform [−0.5, 0.5] distribution. For each factor, observations were assigned to levels uniformly at random so that the the probability of an observation belonging to any specific level was the same for all levels. The diagonal and off-diagonal elements of the random effects covariance matrix for each factor were held fixed across simulation instances and given as 1 and 0.5, respectively.

The spatially varying random terms, E and *b* were generated as images of Gaussian noise, with the appropriate covariance between images induced for *b*. The response images, *Y*, were then calculated using *X, Z, β, b* and E according to equation (1). An isotropic Gaussian filter with a Full Width Half Maximum (FWHM) of 5 was then applied to the response images, *Y*, to induce spatial correlation across voxels. Following this, for each response image, random perturbations were applied to the FSL standard 2mm MNI brain mask in order to generate a corresponding “random mask” image, thus simulating missingness near the edge of the brain (c.f. Supplementary Material Section S6). The “random” masks were applied to the response images to finally obtain a masked smoothed random response, roughly resembling null fMRI data. A visual overview of the data generation process is provided by Figure 2. All NIfTI volumes generated for simulation were of dimension (100 × 100 × 100) voxels. It must be stressed that this process was not designed to simulate realistic fMRI data rigorously, but rather data that had approximately the same shape, size, smoothness, and degree of missingness as might be observed in real fMRI data.

**Figure 2:**
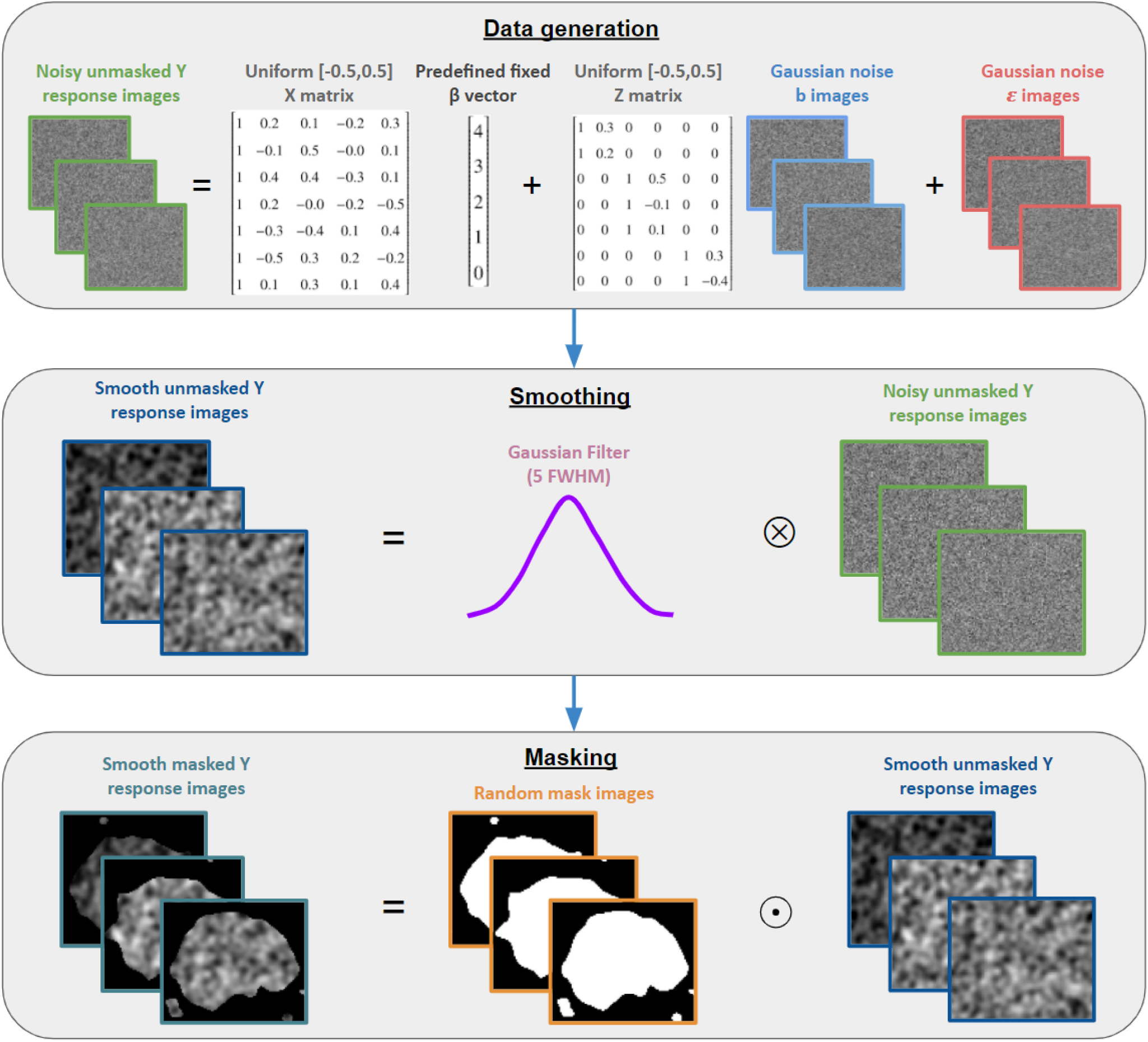
A visual representation of the pipeline employed to generate the simulated data of Section 3.1. The first box depicts the model which was used for data generation, notably highlighting that *ϵ* and *b* varied across space. The second box details the smoothing process, with the ⊗ symbol representing convolution in this instance. The third box details the masking stage, with ⊙ representing the Hadamard (element-wise) product.

In order to assess the accuracy and performance of the parameter estimation in each simulation instance, the R function lmer was also used to obtain parameter estimates for each voxel in the analysis mask. To measure computation speed, we define the ‘Serial Computation Time’ (SCT) for BLMM parameter estimation as the time in seconds that would have been spent executing the FSFS algorithm if the computation performed by each node had been run back-to-back in serial. The serial computation time for lmer parameter estimation is similarly defined as the total amount of time that would have been spent executing the function ‘optimizeLmer’ had all computation been performed in serial. The performance of lmer and BLMM was contrasted in terms of SCT averaged across simulation instances, whilst the parameter estimates produced by lmer and BLMM were compared in terms of image-wide mean absolute difference to assess the correctness of the FSFS algorithm employed by BLMM. It is noted here that the computation of product forms is not included in our evaluation of SCT for BLMM. However, we do not believe this has impacted the results of our analysis as preliminary testing demonstrated that the time spent performing product form computation was many orders of magnitude (approximately 10^5^) smaller than the time spent performing REML estimation using the FSFS algorithm.

The primary purpose in choosing lmer as a baseline for comparison was to demonstrate the substantial computational benefits of using the BLMM pipeline for mass-univariate analysis in the place of naive computation via ‘for loops’ and lmer. We stress that it is not the author’s intention for the results of this analysis to be interpreted as a reflection on the ability of the lme4 package, which is not designed for use in the mass-univariate setting, but rather the efficiency of the FSFS algorithm when combined with vectorised computation. To ensure the missingness capabilities of BLMM were exhaustively tested, each simulated analysis employed a lenient missingness threshold of 50%.

In sum, the simulations we have described assess BLMM’s (i) correctness in terms of parameter estimation, (ii) ability to handle missing data, and (iii) computation speed for parameter estimation. All reported results were obtained using an HPC cluster with Intel(R) Xeon(R) Gold 6126 2.60GHz processors each with 16GB RAM.

### 2.3. Real Data Methods

As a demonstration of the large-scale capabilities of BLMM, here we provide an example involving a much larger model than those considered during the simulations of Section 2.2. In this example, we utilise data from the UK Biobank, in which repeated measures were recorded for 2461 subjects, each of whom completed a “faces vs shapes” task twice across separate visits. In FSL, a first-level analysis was conducted independently for each visit for each subject. In each first-level analysis, the task design was regressed onto Blood Oxygenation Level Dependent (BOLD) response. From each first-level design, a Contrast Parameter Estimate (COPE) map was generated that represented the average difference in the BOLD response between subjects being shown images of faces and images of abstract shapes.

At the group level, BLM and BLMM were used to perform parameter estimation for three models, each designed to estimate the average group level ‘faces>shapes’ response. In each model, the response images, *Y*, were the COPE maps that were output by FSL during the first-level analysis, registered to MNI space. The fixed-effects design matrix, *X*, in each model included an intercept, the cross-sectional effect of age (average age per subject), longitudinal time (inter-visit time in years, centred by subject), sex, a cross-sectional age-sex interaction effect and the Townsend deprivation index (a measure of socio-economic status).

The first of the three group-level analyses (Model 1) was a linear regression model estimated using BLM. As it was a standard linear regression, this model included no random-effects design matrix, *Z*. The second group-level model (Model 2) was analysed using BLMM and included a subject-level random intercept in the random-effects design matrix. The third of the group-level models (Model 3) was run using BLMM and included both a subject-level random intercept and a subject-level random slope for longitudinal time in the random-effects design matrix. For each model, approximate Wald *T* −statistics were computed, using the methods of Section 2.1.4, for the model intercept (the group-level average response to the ‘faces>shapes’ task), the cross-sectional age and longitudinal time. Once the parameters of each model had been estimated, to assess goodness of fit, model comparison was performed using the LRT method detailed in Section 2.1.4.

It must be noted that, as each model contained two observations per subject and model 2 contained two random effects per subject, model 2 contained the same number of observations as random effects. As a result of this, model 2 may be expected to be unidentifiable for many voxels, as parameter estimation for any voxel with missing data will attempt to model at least two random effects using less than two visits. This choice of model is deliberate as model 2 is an extreme use-case that both stress-tests the BLMM code and serves as a clear example of a model that is expected to be rejected by the LRT procedure. It must be stressed that, by including model 2 in this example, it is not our intention to endorse estimation of models that have been ill-specified in this manner. Model 2 is provided purely for the purposes of demonstration and can only be estimated in BLMM by turning off a ‘safeMode’ setting during input specification.

The primary purpose of the analyses described above is to demonstrate BLMM’s usage in practice and highlight BLMM’s efficiency and scalability via a worked example. In order to assess computational efficiency, model 2 was also estimated voxel-wise using lmer and, as in Section 2.2, serial computation times were recorded for parameter estimation for both lmer and BLMM. However, the same comparison could not be performed for model 3, as the default settings in lmer will not allow for models with equal numbers of random effects and observations to be estimated. In Section 3.2, results are reported for both Likelihood Ratio Tests (model 1 vs model 2 and model 2 vs model 3), with *p*-value significance maps for the approximate Wald *T* -tests then being provided for the selected model. All reported significance regions were obtained using a 5% Bonferonni corrected threshold. All analysis results were obtained using a SGE cluster with Intel(R) Xeon(R) Gold 61262.60GHz processors, each with 16GB RAM.

## 3. Results

### 3.1. Simulation Results

Across the nine simulation settings outlined in Section 3.1 (three designs across three sample sizes), all parameter estimates and maximised restricted likelihood criteria produced by BLMM were near identical to those produced by lmer. In particular, extremely strong agreement was observed between the parameter estimates produced by BLMM and lmer for both voxels with missing observations and voxels with all observations present, illustrating BLMM’s capacity for handling missing data. For each experimental design, the largest mean absolute difference for parameter estimation was observed in the estimation of the random effects covariance parameters (the unique non-zero elements of *D*) in the *n* = 200 setting. The observed absolute mean difference in the random effects covariance parameter estimates produced by BLMM and lmer for this setting were 6.86 × 10^−9^, 4.39 × 10^−5^ and 6.02 × 10^−3^ for designs 1, 2 and 3 respectively. Assuming double precision and a machine level tolerance of 2^−26^ ≈ 1.49 × 10^−8^, it can be seen that this means the estimates produced by BLMM and lmer were indistinguishable at the machine precision level for design 1 and very similar for designs 2 and 3. Similar results may also be observed across simulations for the maximised REML criterions produced by BLMM and lmer (see Supplementary Material Sections S7-S10).

The observed SCTs for each simulated setting are presented in Fig. 3. As can be seen from Fig. 3, BLMM significantly outperformed computation via ‘for loops’ and lmer and, notably, appeared to maintain an approximately constant computation time as the number of observations, *n*, increased. In contrast, computation time for lmer appeared to increase as the number of observations increased. These results match expectation as, as noted in Section 2.1.2, the BLMM pipeline begins by computing the product forms and discarding any model matrices which had dimensions scaling with *n*. This means that during parameter estimation (the most intensive stage of the BLMM pipeline, which dominates the computation time) it is expected that computation should not scale as a function of *n*. As computation time does scale as a function of *q*, however, it should be noted that, as shown in Fig. 3, when *q* and *n* are close in magnitude the benefits of the product form approach to computation are less pronounced.

**Figure 3:**
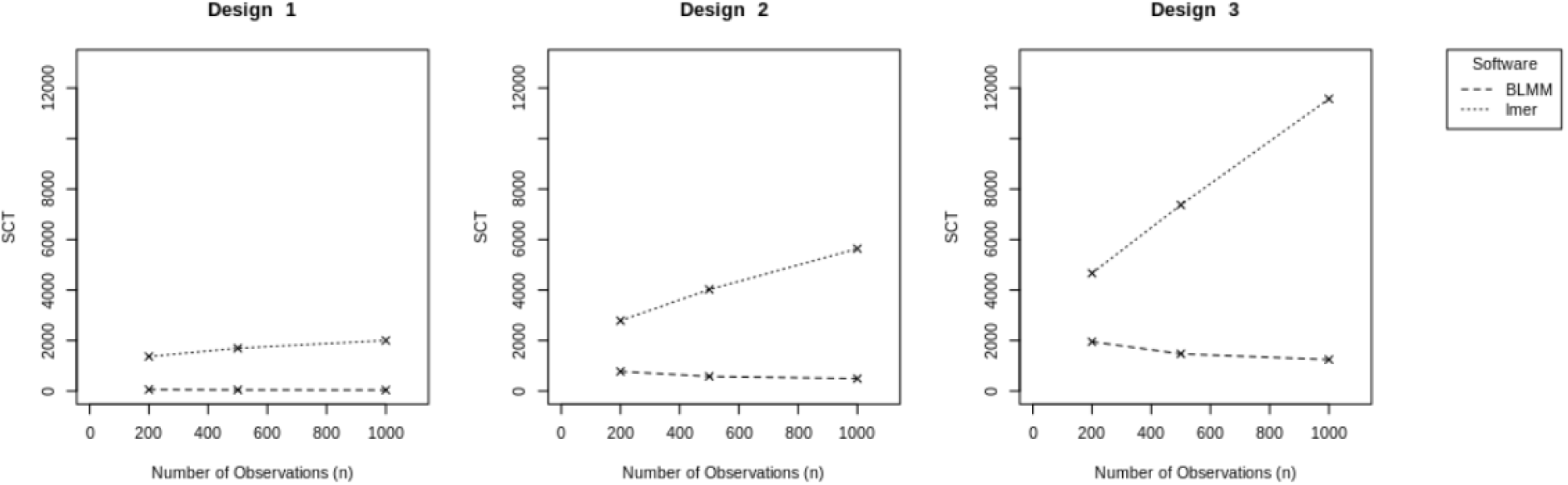
Observed serial computation times for each experimental design, displayed as a function of the number of observations, *n*. Displayed are the SCT in seconds for BLMM (dashed) and lmer (dotted).

The results of Fig. 3 demonstrate BLMM’s strong computational efficiency for analysing the designs which were simulated. However, it should be noted that the difference in SCT between lmer and BLMM may be smaller than those observed in these simulations for analyses involving designs in which the second dimension of the random effects design matrix, *q*, is very large. The reason for this is that, as *q* increases, the storage requirements associated with each voxel increase and, as a result, parameter estimation can be performed for fewer voxels concurrently via vectorisation. In the simulations presented in this section, *q* was set to 100 in designs 1 and 2 and set to 50 in design 3. We note that it would be of benefit to conduct further detailed analysis of BLMM’s performance compared to that of lmer for models with higher values of *q*. However, at present, such comparative simulations are practically infeasible due to the inordinate computation time they would require (the reported simulations required several months to run, generated over five million simulated brain images and executed approximately 10^7^ univariate LMM analyses in lmer). Whilst such extensive simulations may not be possible, we highlight here that the analyses presented in Section 3.2 provide further evidence of the BLMM’s strong performance for two models in which *q* is much larger than considered in the simulations presented in this section (*q* = 2461 and *q* = 4922 in models 1 and 2, respectively).

### 3.2. Real Data Results

The results of the UK Biobank real data example of Section 2.3 are given by Fig. 4. The first and second rows of Fig. 4 display the regions of significance obtained from LRTs for model comparison between models 1 and 2, and models 2 and 3, respectively. The results of the first LRT, comparing model 2 (the random intercept model) and model 1 (the linear regression), highlight most of occipital, parietal and frontal lobes. This observation suggests that, in these regions, evidence was observed that the inclusion of subject-level random intercepts substantially influenced the outcome of the analysis. This conclusion is reasonable and reflects regions where there is non-negligible BOLD response to this task.

**Figure 4:**
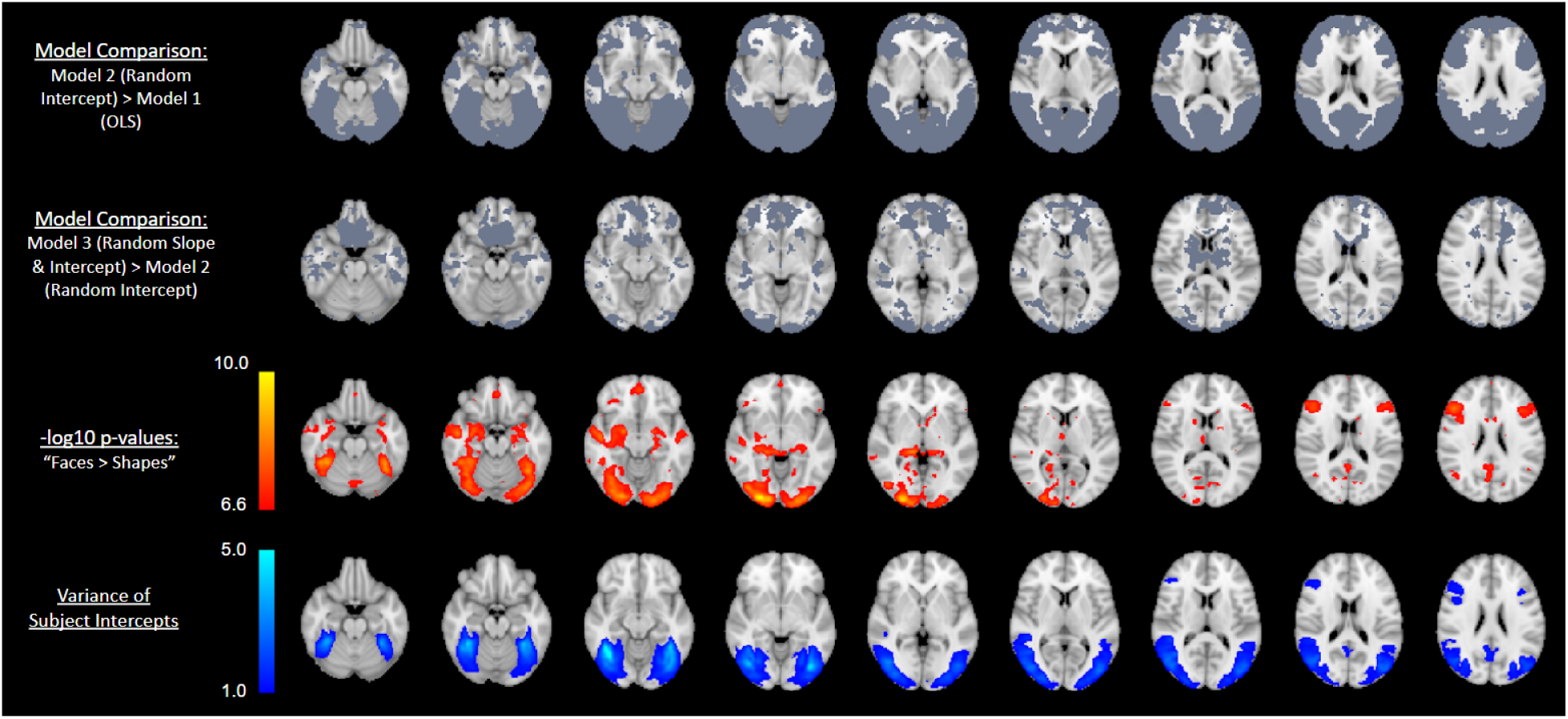
First row: LRT results for comparison of model 1 and model 2; for this row, regions highlighted demonstrate where evidence was found that inclusion of a random subject intercept significantly affected the results of the analysis. Second row: LRT results for comparison of model 2 and model 3; regions highlighted indicate where the inclusion of a random slope for longitudinal time significantly impacted the analysis. Third row: Effect of “Faces > Shapes” p-values, given on the − log_10_(*p*) scale, are shown; for this row, voxels shown are those Bonferroni-significant for the “Faces > Shapes” contrast. Fourth row: Variance estimates for the subject-level random intercept, thresholded at 1. Voxels shown are those which displayed notable variability at the subject-level for the random intercept included in the model.

The second LRT, comparing model 3 (the random intercept and slope model) to model 2, only identifies two different regions: orbitofrontal areas, which are often subject to signal loss, and white matter areas in the centre of brain, which have the poorest SNR with this multicoil acquisition. The sporadic and noisy appearance of the regions identified here could indicate that these regions were influenced by idiosyncratic temporal changes due to variation in head placement or simply by large random temporal changes. In either case, it can be seen that the inclusion of a random slope in the analysis had little impact on anatomical regions of practical relevance to the ‘faces vs shapes’ task. The conclusion of the LRTs is, therefore, that model 2 should be chosen as the suitable model upon which inference can be performed using approximate Wald *T* -tests.

Of the three contrasts that were estimated for model 2 using the approximate Wald *T* -test, only the first contrast (the main effect of the “Faces vs Shapes” task) reported significant regions following a 5% Bonferroni-corrected threshold. The thresholded p-value significance map for this contrast, reported on the − log_10_(*p*) scale, is displayed on the third row of Fig. 4. In this instance, the occipital lobe, known for its role in processing perceptual information (c.f. Rehman and Khalili (2019)), has been identified as significant. This result matches expectation, given the visual nature of the faces vs shapes task.

Also provided in the fourth row of Fig. 4 is an image representing the estimated subject-level variability for the random intercept. This image serves as a useful diagnostic, as it highlights regions at which inclusion of the random intercept notably contributed to the analysis results. In this example, it can be seen that the regions at which the subject-level variance estimate exceeded 1 exhibit similarity to those regions which were deemed significant during the approximate Wald *T* -test. This illustrates the vital role that the inclusion of a random intercept played in obtaining the results of this analysis and reinforces the conclusion of the LRT testing procedure.

In total, computation time in BLMM for this analysis took approximately 55 minutes for model 2 and 4 hours for model 3, using 500 nodes. The extreme computation time observed for model 3 can be attributed to the identifiability problems noted in Section 2.3, with parameter estimation for approximately 150 voxels exceeding the default maximum iteration limit. By way of comparison, for model 2, the SCT observed for BLMM was approximately 77.6% of that observed for lmer. As noted previously, this difference is less pronounced than those observed in the simulations of Section 3.1, as the second dimension of the random effects design matrix is extremely large (*q* = 2461) and close in magnitude to the value of *n* (*n* = 4922). The linear regression model, model 1, was run using BLM and took approximately 5 minutes using 60 computational nodes. The full analysis results for all three models are publicly available and may be accessed on NeuroVault (see the data and code availability declaration).

## 4. Discussion and Conclusion

In this work, we have detailed and presented BLMM, a freely available software package for performing LMM parameter estimation and inference on large-*n* fMRI datasets. LMM computation for fMRI is an extremely computationally intensive task and, as a result, the work presented in this paper is both informed and limited by the currently available support in terms of software and technology. For this reason, BLMM is a continually evolving project, and it is expected that much of the methodology currently implemented in BLMM may gradually change over time.

A large driving force motivating the approaches taken in this work is the current lack of available support for broadcasted sparse matrix operations. Whilst the available support for sparse matrix methodology in programming languages such as MatLab and Python has substantially improved in recent years, currently, there is little support for performing sparse matrix operations concurrently many times. Unfortunately, this substantially limits the utility of sparsity-based approaches to LMM estimation in the context of fMRI analysis. This is due to the substantial over-heads which would be accrued if LMM estimation were performed independently for each voxel in an image. An undesirable ramification of this is that the BLMM code, which has been explicitly optimised to account for the patterns of sparsity present in single-factor models (see Supplementary Material Section S4), may not provide comparable performance for multi-factor models in which *q* is very large. However, as programming languages evolve and new support becomes available, this is expected to change, and we predict that future development of BLMM is highly likely to incorporate sparse matrix methodology into its approach to analysing multi-factor models. For this reason, we suggest that the incorporation of sparse matrix methodology into BLMM may form a substantial basis for future development.

Another potential avenue for future development focuses on how BLMM currently handles file input and output on HPC clusters. In recent years, in Python especially, there has been a strong movement towards standardising and streamlining cluster-based computation. A project of particular note is the Dask Python package, which acts as a standardised specification to encode parallel algorithms and file I/O (Rocklin (2015)). As noted in Section 1.1.1, Dask is heavily employed by the AFNI package 3dLME and provides substantial benefits both in terms of computation time and ease of distribution. Although many of the broadcasted operations employed by BLMM are yet to be supported by Dask, we suggest that incorporating the approach of AFNI’s 3dLME by integrating Dask with the existing BLMM code-base may provide further speed improvements and could form the foundation of future development within BLMM.

There is also much room for further theoretical development of BLMM. In our previous work, Maullin-Sapey and Nichols (2021), we presented a general approach for performing constrained optimisation to enforce structure in the random effects covariance matrix. This concept may be of utility in a wide range of commonplace applications, as structured covariance matrices are a feature of many popular statistical models. Such applications include, for example, modelling genetic components in twin studies and auto-correlation in time-series analyses. Whilst the approaches outlined in our previous work are sufficiently general to perform such analyses in the univariate setting, it is not immediately apparent whether they may be feasibly executed on a mass-univariate scale. For this reason, we suggest that the potential for enforcing covariance structure in BLMM merits further investigation.

Another potential direction for future work stems from the observation that the BLMM framework could be used to support remote distributed computation when raw data cannot be centrally stored. Such prohibitive situations are common when, for example, the raw data is extremely large, or subject to privacy constraints preventing it from being shared. The BLMM framework may be of utility in such situations as each remote site may compute their product forms using only their data and their portion of the design matrix, apply some form of differential privacy protection to the results (e.g. add calculated amounts of noise to the product forms to preserve privacy), and send the results to a central coordinate that never sees the raw data. Similar approaches to distributed computation have been notably adopted by the Collaborative Informatics and Neuroimaging Suite Toolkit for Anonymous Computation (COINSTAC, c.f. Plis et al. (2016)) and suggest new alternative applications for the BLMM framework.

## 5. Declarations

### 5.1. Data/code availability statement

We have used data from The UK Biobank. The BLMM toolbox, as well as the code used for the simulations and timing comparisons described in Sections 2.2 and 2.3, are available at:

> https://github.com/TomMaullin/BLMM

The BLM toolbox, which is also referenced several times throughout this work, may be found at:

> https://github.com/TomMaullin/BLM

The images generated by BLMM for the three models discussed in Sections 2.3 and 3.2 may be found at:

> Model 1: https://identifiers.org/neurovault.collection:9881
>
> Model 2: https://identifiers.org/neurovault.collection:10448
>
> Model 3: https://identifiers.org/neurovault.collection:10450

The results of the LRT’s for model comparison discussed in Sections 2.3 and 3.2 may also be found at:

> https://identifiers.org/neurovault.collection:10451

### 5.2. Ethics statement

The data we have used was provided by the UK Biobank. The UK Biobank has generic Research Tissue Bank (RTB) approval, which covers our research using the resource.

### 5.5. CRediT authorship contribution statement

Thomas Maullin-Sapey: Conceptualization, Formal analysis, Investigation, Methodology, Software, Visualization, Roles/Writing - original draft, Writing - review & editing. Thomas Nichols: Conceptualization, Funding acquisition, Supervision, Writing - review & editing.

