## Supplementary Material for "BLMM: Parallelised Computing for Big Linear Mixed Models"

### S1 Illustrative Example of BLMM Setup

To illustrate the construction described in Section 2.1.1 of the main text in practice, we provide a brief mock example consisting of a "small" longitudinal group-level analysis. The analysis design contains 3 subjects, each of which has had multiple image acquisitions taken irregularly across repeated visits and has recorded observations for "sex", "age", and "BMI". Only one grouping factor for the random effects is included in the model, the factor "subject", and acquisitions for the first, second and third subjects were taken across 2, 3 and 2 visits, respectively. (It is emphasized that the number of subjects and observations used in this example are far too small to draw meaningful inference from in practice and that this example has only been constructed for illustrative purposes). For this example, if the user wished to model a between-subject random intercept and a between-subject random slope for age using BLMM, the input for the matrices $X, g_1$ and $z_1$, and the full construction of $Z$, may appear as follows:

$$X = \begin{bmatrix} 1 & 1 & 19 & 18.7 \\ 1 & 1 & 21 & 19.2 \\ 1 & 1 & 18 & 24.8 \\ 1 & 1 & 20 & 25.1 \\ 1 & 1 & 24 & 24.6 \\ 1 & 0 & 21 & 20.9 \\ 1 & 0 & 22 & 20.1 \end{bmatrix}, \quad g_1 = \begin{bmatrix} 1 \\ 1 \\ 2 \\ 2 \\ 2 \\ 3 \\ 3 \end{bmatrix}, \quad z_1 = \begin{bmatrix} 1 & 19 \\ 1 & 21 \\ 1 & 18 \\ 1 & 20 \\ 1 & 24 \\ 1 & 21 \\ 1 & 22 \end{bmatrix}, \quad Z = \begin{bmatrix} 1 & 19 & 0 & 0 & 0 & 0 \\ 1 & 21 & 0 & 0 & 0 & 0 \\ 0 & 0 & 1 & 18 & 0 & 0 \\ 0 & 0 & 1 & 20 & 0 & 0 \\ 0 & 0 & 1 & 24 & 0 & 0 \\ 0 & 0 & 0 & 0 & 1 & 21 \\ 0 & 0 & 0 & 0 & 1 & 22 \end{bmatrix}$$

where the four columns of $X$ correspond to an intercept, sex, age and BMI respectively, $g_1$ indicates which observations correspond to which subjects, and $z_1$ includes an intercept and age. In this example, the input for the response vector, $Y$, would be a text file containing a list of the filenames of seven NIfTI images corresponding to each visit across all subjects. During input specification, the user may also specify hypothesis tests to be conducted. For example, to perform an approximate Wald T-test testing for the non-zero effect of BMI in the above example, the user may specify $T$ as the statistic type and $[0,0,0,1]$ as the contrast vector.

### S2 BLM: Parallelised Computing for Big Linear Models

In Section 2.1.2, a "product form"-based method for performing parameter estimation and inference for the Linear Model was referenced. This method served as the initial motivation for BLMM's approach to parallelized parameter estimation and inference for the Linear Mixed Model and is implemented as a sister-project to BLMM under the acronym BLM (Big Linear Models). In this appendix, we give a brief outline of the computational stages of BLM, highlighting the similarity between the approaches of BLM and BLMM.

Much like BLMM, BLM is designed to account for missingness observed as a result of mask-variability. As such, BLM assumes a model setup that is analogous to that described in Section 1.1.2 given by;

$$Y_v = M_v X\beta_v + \varepsilon_v, \qquad \varepsilon_v \sim N(0, \sigma_v^2 M_v)$$

where $M_v$ is the $(n \times n)$-dimensional 'missingness' matrix described in Section 1.1.2 and the missing values in $Y_v$ are encoded as zero. Mirroring the approach of Section 2.1.2, to reduce memory consumption, computation in BLM begins by evaluating the below product forms for each voxel in the analysis mask;

$$P_v = X_v' X_v, \;\; Q_v = X_v' Y_v, \;\; S_v = Y_v' Y_v,$$

where $X_v = M_v X$. Following computation of $P_v, Q_v$ and $S_v$, the matrices $X_v, Y_v$ and $Z_v$ are redundant and discarded from memory. By employing vectorized computation in a manner similar to that described in Section 2.1.3, BLM then evaluates the ordinary least squares estimators using the product forms as follows:

$$\hat{\beta}_v = P_v^{-1} Q_v, \qquad \hat{\sigma}_v^2 = S_v - 2Q_v'\hat{\beta}_v + \hat{\beta}_v' P_v \hat{\beta}_v$$

Finally, in a similar manner to that described in Section 2.1.4, BLM supports null-hypothesis testing via Wald statistics. Neglecting the voxel subscript $v$, Wald $T-$ and $F-$statistics are calculated using the product forms as follows:

$$T = \frac{L\hat{\beta}}{\sqrt{\hat{\sigma}_v^2 L P^{-1} L'}}, \qquad F = \frac{\hat{\beta}' L' (L P^{-1} L')^{-1} L\hat{\beta}}{\hat{\sigma}^2 \text{rank}(L)}$$

where $L$ is a fixed and known contrast vector. For further information on the BLM toolbox, see:

https://github.com/TomMaullin/BLM

### S3 Computational efficiency for product form evaluation

In Section 2.1.2, pseudocode demonstrates how BLMM evaluates the product forms in a computationally efficient manner. For clarity and conciseness, however, several practical details were neglected from Algorithm 1. Therefore, in this section, to complement the discussion of Section 2.1.2, we provide a brief overview of several practical implementation details which were neglected from Algorithm 1.

Firstly, we highlight that in Algorithm 1, several operations were conceptualized as involving the missingness matrix $M_v^{(b)}$ which has dimension $(\frac{n}{B} \times \frac{n}{B})$. Whilst the use of the matrix $M_v^{(b)}$ in this situation is theoretically justified, construction of such a matrix is computationally infeasible due to the large amount of memory required in order to store $M_v^{(b)}$ for all voxels. Here we note that, in practice, construction of the matrix $M_v^{(b)}$ is unnecessary. Instead, multiplications such as $M_v^{(b)}X_v^{(b)}$ can be evaluated incrementally by replacing rows of $X_v^{(b)}$ with zeros in an appropriate fashion.

Secondly, as a result of between-voxel design commonality (c.f. Section 1.2.2), we highlight that typically $P_v$, $R_v$ and $U_v$ will take the same values for many voxels. As such, $P_v$, $R_v$, and $U_v$ do not need to be computed and stored separately for every voxel. Although not noted in Algorithm 1, BLMM utilises this between-voxel commonality during the product form stage of computation to reduce storage costs by storing only the unique instances of $P_v$, $R_v$ and $U_v$. As it is often the case that there are large contiguous regions of the analysis mask over which there is little missingness, storing $P_v$, $R_v$ and $U_v$ in this manner is expected to be much more memory efficient than storing them for each voxel individually.

Thirdly, in Algorithm 1, it appears there are many "for loops" which are being serially executed across voxels. Here we highlight that the operations inside these loops utilise only conceptually simplistic operations such as matrix multiplication and addition. Consequently, in practice, these loops can be realised in a quick, efficient parallelised manner by using vectorised computation (c.f. Section 2.1.3). It must also be noted, however, that in cases when $R_v$ and $U_v$ are particularly large, on each node, BLMM may not be able to evaluate $R_v$ and $U_v$ for all voxels concurrently due to the high RAM usage that would be required. In such cases, BLMM will split $\{X_v^{(b)}\}$ and $\{Z_v^{(b)}\}$ voxel-wise into further batches and, on each node, consider each batch in serial.

Finally, it is also noted that for models which contain only one random factor, $U_v$ becomes block diagonal. In such situations, substantial gains in computation time and memory consumption may be made via careful consideration of the structure of $U_v$. The benefits of this approach are most extreme when the model design contains only one random factor and one random effect. In this case, $U_v$ is diagonal, and BLMM needs only to evaluate and store the diagonal elements of $U_v$. Such considerations are discussed in greater detail in the following section, Section S4.

#### S4 Computational efficiency for single-factor designs

This section outlines two common scenarios in which structural features of the random effects covariance and design matrices, $D$ and $Z$, allow for further improvements in terms of computational efficiency in the BLMM pipeline. To illustrate how BLMM exploits the structure of $D$ and $Z$ to achieve improved computational performance in each of these scenarios, we shall use Equation (8) as an informative example throughout this section. For the remaining expressions listed in Sections 2.1.1-2.1.4, we note that similar adjustments are possible but, for brevity, will not be listed here.

The first scenario in which BLMM accounts for the structure of $Z$ and $D$ computationally is the setting in which the experimental design contains a single random factor which groups multiple random effects (i.e. $r = 1$ and $q_1 > 1$). In such instances, $U$ (which is equal to $Z'Z$) and $D$ are both block diagonal, with $D$ consisting of only one block, $D^1$, which is of dimension $(q_1 \times q_1)$, repeated $l_1$ times along it's diagonal (i.e. $D = I_{l_1} \otimes D^1$). By noting the structure of $D$ and $U$, and neglecting the subscript $s$ for iteration number, Equation (8) may be rewritten in this setting as follows:

$$\begin{aligned} X'V^{-1}X &= P - \sum_{j=1}^{l_1} R_{(j)} D^1 (I + D^1 U_{(j)})^{-1} R'_{[j]} \\ Z'V^{-1}Z &= \bigoplus_{j=1}^{l_1} \left( U_{(j)} - U_{(j)} D^1 (I_{q_1} + D^1 U_{(j)})^{-1} U_{(j)} \right) \\ Z'V^{-1}e &= T' - R'\beta - \bigoplus_{j=1}^{l_1} \left( U_{(j)} D^1 (I_{q_1} + D^1 U_{(j)})^{-1} \right) (T' - R'\beta) \end{aligned} \tag{S1}$$

where $U_{(j)}$ and $R_{(j)}$ are shorthand for $Z'_{(1,j)}Z_{(1,j)}$ and $X'Z_{(1,j)}$ respectively, and $\oplus$ is the direct sum. Despite appearing notationally convoluted, the above expressions are much more convenient than Equation (8) for computational purposes. This is due to the notably smaller dimensions of the matrices involved in the right hand side of Equation (S1) in comparison to those employed in Equation (8).

For example, consider the dimensions of the matrices which are inverted in Equation (S1), $\{(I_{q_1} + D^1 U_{(j)})\}_{j \in \{1,\dots,l_1\}}$, and the matrix which is inverted in Equation (8), $(I_q + DU)$. The former matrices have dimension $(q_1 \times q_1)$ whilst the latter has dimension $(q \times q)$. To illustrate the significance of this observation, consider a setting in which 2 random effects are modelled across 100 levels (subjects or time-points, for example). In such a scenario, $q = 200$ whilst $q_1 = 2$ and, by using the above reformulation of Equation (8), the inversion of a matrix of dimension $(200 \times 200)$ can be substituted for the much faster operation of inverting 100 matrices of dimension $(2 \times 2)$. When applied in the mass-univariate setting, such improvements in efficiency can reduce computation time from hours to seconds.

The second scenario to which computation in the BLMM pipeline has been tailored is the setting in which the experimental design contains a single random factor that groups a single random effect (i.e. $r = 1$ and $q_1 = 1$). In this instance, the matrices $D$ and $U$ are diagonal, with $D$ containing a single scalar value, $d$, repeated $l_1$ times along the diagonal. Consequently, by denoting $u$ as the vector composed of the diagonal elements of $U$, Equation (8) can be rewritten in this setting as follows:

$$\begin{aligned} X'V^{-1}X &= P - d(RR' \odot \text{diag}(1 \oslash (1 + du)) \\ Z'V^{-1}Z &= \text{diag}(u \odot (1 - du) \oslash (1 + du)) \\ Z'V^{-1}e &= (1 - du) \oslash (1 + du) \odot (T' - R'\beta) \end{aligned} \tag{S2}$$

where $\odot$ and $\oslash$ represent Hadamard (element-wise) multiplication and division, respectively, and diag is the unique function which maps an arbitrary $(m \times 1)$ vector, $x$, to an $(m \times m)$ diagonal matrix with the elements of $x$ listed along its diagonal. As in the previous scenario, in which the use of Equation (S1) greatly reduced the time complexity of computation, the use of Equation (S2) can also provide strong improvements in terms of computational speed. A notable example of this can be seen by contrasting the time complexity of the evaluation of $Z'V^{-1}Z$ obtained using the right-hand side of Equation (S2) against that obtained using Equation (8). The time complexity of the former is $O(q)$, whereas the time complexity of the latter is greater than $O(q^2)$. As a consequence, in this scenario, the matrix

$Z'V^{-1}Z$, which is utilised many times throughout the BLMM pipeline, may be evaluated in a much faster manner via the use of Equation (S2) than by use of Equation (8).

In each of the above scenarios, it must further be noted that, by utilising the internal structure of $Z$, memory consumption can also be significantly reduced. For example, when storing the product form $U = Z'Z$, only the non-zero entries of $U$ must be recorded. This reduces the storage cost associated to $U$ from a complexity of $O(q^2)$ to; $O(q_1 q)$ in the scenario in which $U$ is block-diagonal, and $O(q)$ in the scenario in which $U$ is diagonal. We note here that sparse matrix methods could theoretically also be employed to obtain further improvements in computational time and storage. However, due to the present lack of vectorised support available for sparse matrix operations, such improvements are currently infeasible in practice for mass-univariate analyses. For this reason, BLMM currently does not utilise sparse matrix methodology.

#### S5 Expressions for degrees of freedom estimation

In Section 2.1.4, a general overview of BLMM's approach to degrees of freedom estimation was provided, with several expressions being omitted for the purposes of concision. In this section, we list the expressions which were omitted from Section 2.1.4. To be precise, here we provide closed-form expressions for the unknown covariance matrix $\text{Var}(\hat{\eta})$, and the gradient vector $\frac{dS^2(\hat{\eta})}{d\hat{\eta}}$. To state these, we partition $\text{Var}(\hat{\eta})$ and $\frac{dS^2(\hat{\eta})}{d\hat{\eta}}$ block-wise as follows:

$$\text{Var}(\hat{\eta}) = \begin{bmatrix} \mathscr{I}_{\hat{\sigma}^2,\hat{\sigma}^2} & \mathscr{I}_{\hat{\sigma}^2,\hat{D}^1} & \cdots & \mathscr{I}_{\sigma^2,\hat{D}^r} \\ \mathscr{I}_{\hat{D}^1,\hat{\sigma}^2} & \mathscr{I}_{\hat{D}^1,\hat{D}^1} & \cdots & \mathscr{I}_{\hat{D}^1,\hat{D}^r} \\ \vdots & \vdots & \ddots & \vdots \\ \mathscr{I}_{\hat{D}^r,\hat{\sigma}^2} & \mathscr{I}_{\hat{D}^r,\hat{D}^1} & \cdots & \mathscr{I}_{\hat{D}^r,\hat{D}^r} \end{bmatrix}, \qquad \frac{dS^2(\hat{\eta})}{d\hat{\eta}} = \begin{bmatrix} \frac{dS^2(\hat{\eta})}{d\hat{\sigma}^2} \\ \frac{dS^2(\hat{\eta})}{d\text{vec}(\hat{D}^1)} \\ \vdots \\ \frac{dS^2(\hat{\eta})}{d\text{vec}(\hat{D}^r)} \end{bmatrix}.$$

where $\mathscr{I}_{A,B}$ is shorthand for the covariance matrix $\text{Cov}(\frac{dl(\hat{\theta})}{d\text{vec}(A)}, \frac{dl(\hat{\theta})}{d\text{vec}(B)})$. Given the above partition $\text{Var}(\hat{\eta})$ may now be stated block-wise as follows:

$$\mathscr{I}_{\hat{\sigma}^2,\hat{\sigma}^2} = \frac{n}{2}\hat{\sigma}^{-4},$$

$$\text{for } k \in \{1,...r\}, \qquad \mathscr{I}_{\hat{\sigma}^2,\hat{D}^k} = \frac{1}{2\hat{\sigma}^2}\text{vec}'\bigg(\sum_{j=1}^{l_k} Z'_{(k,j)}\hat{V}^{-1}Z_{(k,j)}\bigg),$$

$$\text{for } k_1,k_2 \in \{1,...r\}, \qquad \mathscr{I}_{D^{k_1},D^{k_2}} = \frac{1}{2}\sum_{j=1}^{l_{k_2}}\sum_{i=1}^{l_{k_1}}\bigg(Z'_{(k_1,i)}V^{-1}Z_{(k_2,j)} \otimes Z'_{(k_1,i)}V^{-1}Z_{(k_2,j)}\bigg),$$

and the gradient vector, $\frac{dS^2(\hat{\eta})}{d\hat{\eta}}$, may be stated block-wise as:

$$\frac{dS^2(\hat{\eta})}{d\hat{\sigma}^2} = L(X'\hat{V}^{-1}X)^{-1}L',$$

$$\text{for } k \in \{1,...r\}, \qquad \frac{dS^2(\hat{\eta})}{d\text{vec}(\hat{D}^k)} = \hat{\sigma}^2\bigg(\sum_{j=1}^{l_k}\hat{B}_{(k,j)} \otimes \hat{B}_{(k,j)}\bigg),$$

where $\hat{B}_{(k,j)}$ is given by $\hat{B}_{(k,j)} = Z'_{(k,j)}\hat{V}^{-1}X(X'\hat{V}^{-1}X)^{-1}L'$ and $\hat{V} = I_n + Z\hat{D}Z'$. For use in the BLMM pipeline, the above expressions may be reformulated to be given in terms of the product forms of Section 2.1.2 via use of Equation (8). The reformulated expressions are lengthy, however, and therefore will not be stated in full here. Further discussion of the above expressions can be found in our previous work, Maullin-Sapey and Nichols [2021].

#### S6 Random mask generation

In this section, we detail the process used to generate the random masks which were applied to the simulated data of Section 2.2. To create randomly deformed masks for the simulated analyses, the standard T1 2mm MNI152 brain mask, which is available in the FSL software package, was employed. Mask generation began by first eroding the standard T1 2mm mask by approximately three voxels. This was achieved by smoothing the binary mask with an isotropic Gaussian kernel of 4 Full Width Half Maximum (FWHM) and then re-thresholding the resultant smooth image at 0.7. The purpose of this erosion was to create a mask that was approximately the same shape and size as the standard T1 2mm MNI152 brain mask but with some voxels missing near cortical boundaries and the edge of the brain. Following this, random deformations in the size and shape of the eroded analysis mask were generated. This process was achieved by multiplying an image of $N(0,1)$ Gaussian noise by 8, adding the result to the eroded T1 2mm mask and smoothing the resultant sum with an isotropic Gaussian kernel of 10 FWHM. The final image was then re-thresholded at 0.6. This process was established empirically to ensure that the masks produced had approximately the same quantity of voxels as a standard fMRI analysis mask, with enough random deformation present to exhaustively test the missingness handling capacity of BLMM.

Using the masks generated by this process, the final analysis mask in each simulation instance (following the application of a 50% missingness threshold, c.f. Section 2.1.1) contained on average 217930.6 voxels, with a standard error of approximately 127.0 voxels across simulation instances. This meant that the final analysis mask occupied approximately 21.8% of each NIfTI volume. As noted earlier, this is, by design, similar to the 22.8% of the NIfTI volume that is occupied by the original FSL T1 2mm brain mask and may be expected to be occupied in a standard fMRI analysis. On average, in each simulation instance, approximately 87483.9 voxels had missing data (40.1% of the total number of voxels in the analysis mask), and 130446.7 voxels had full observations across all volumes generated (59.9%). We emphasize here that this extreme degree of subject-mask variability was deliberately simulated to stress test the missingness handling and time efficiency capabilities of BLMM.

#### S7 Mean Absolute Difference for Parameter Estimation (n=200)

| Method | Missing-data voxels | Full-data voxels | All voxels |
|---|---|---|---|
| *Simulation* 1 | | | |
| $\beta$ | $6.86 \times 10^{-9}$ | $4.75 \times 10^{-9}$ | $5.45 \times 10^{-9}$ |
| | $(1.32 \times 10^{-11})$ | $(9.14 \times 10^{-12})$ | $(1.02 \times 10^{-11})$ |
| $\sigma^2$ | $1.43 \times 10^{-9}$ | $1.06 \times 10^{-9}$ | $1.18 \times 10^{-9}$ |
| | $(3.10 \times 10^{-12})$ | $(2.21 \times 10^{-12})$ | $(2.46 \times 10^{-12})$ |
| $D$ | $8.26 \times 10^{-6}$ | $5.20 \times 10^{-6}$ | $6.20 \times 10^{-6}$ |
| | $(1.67 \times 10^{-8})$ | $(1.10 \times 10^{-8})$ | $(1.24 \times 10^{-8})$ |
| *Simulation* 2 | | | |
| $\beta$ | $4.39 \times 10^{-5}$ | $2.34 \times 10^{-5}$ | $3.05 \times 10^{-5}$ |
| | $(2.05 \times 10^{-7})$ | $(1.43 \times 10^{-7})$ | $(1.62 \times 10^{-7})$ |
| $\sigma^2$ | $6.12 \times 10^{-6}$ | $3.33 \times 10^{-6}$ | $4.29 \times 10^{-6}$ |
| | $(2.44 \times 10^{-8})$ | $(1.70 \times 10^{-8})$ | $(1.92 \times 10^{-8})$ |
| $D$ | $2.54 \times 10^{-4}$ | $1.40 \times 10^{-4}$ | $1.79 \times 10^{-4}$ |
| | $(1.24 \times 10^{-5})$ | $(8.59 \times 10^{-6})$ | $(9.78 \times 10^{-6})$ |
| *Simulation* 3 | | | |
| $\beta$ | $1.70 \times 10^{-5}$ | $8.01 \times 10^{-6}$ | $1.11 \times 10^{-5}$ |
| | $(1.13 \times 10^{-7})$ | $(7.53 \times 10^{-8})$ | $(8.73 \times 10^{-8})$ |
| $\sigma^2$ | $1.27 \times 10^{-6}$ | $5.44 \times 10^{-7}$ | $7.95 \times 10^{-7}$ |
| | $(7.13 \times 10^{-9})$ | $(4.11 \times 10^{-9})$ | $(5.09 \times 10^{-9})$ |
| $D$ | $6.02 \times 10^{-3}$ | $2.93 \times 10^{-3}$ | $3.99 \times 10^{-3}$ |
| | $(4.10 \times 10^{-5})$ | $(2.72 \times 10^{-5})$ | $(3.17 \times 10^{-5})$ |

Table S1: Mean absolute difference in the estimates produced by lmer and BLMM for $\beta, \sigma^2$ and $D$, averaged across voxels and simulation instances. Each reported average is an image-wide mean, taken across approximately $218,000$ voxels, further averaged across 1000 simulation instances. In each simulation instance, the model employed included 200 observations generated according to the methods outlined in Section 2.2. Empirical standard errors, taken across simulation instances, are also given in brackets underneath each entry in the table.

#### S8 Mean Absolute Difference for Parameter Estimation (n=500)

| Method | Missing-data voxels | Full-data voxels | All voxels |
|---|---|---|---|
| *Simulation* 1 | | | |
| $\beta$ | $1.54\times10^{-9}$ | $1.09\times10^{-9}$ | $1.26\times10^{-9}$ |
| | $(1.96\times10^{-12})$ | $(1.52\times10^{-12})$ | $(1.60\times10^{-12})$ |
| $\sigma^2$ | $3.48\times10^{-10}$ | $2.54\times10^{-10}$ | $2.89\times10^{-10}$ |
| | $(2.51\times10^{-13})$ | $(1.69\times10^{-13})$ | $(1.78\times10^{-13})$ |
| $D$ | $1.13\times10^{-6}$ | $6.64\times10^{-7}$ | $8.40\times10^{-7}$ |
| | $(1.26\times10^{-9})$ | $(8.30\times10^{-10})$ | $(9.06\times10^{-10})$ |
| *Simulation* 2 | | | |
| $\beta$ | $2.00\times10^{-6}$ | $6.63\times10^{-7}$ | $1.17\times10^{-6}$ |
| | $(1.13\times10^{-8})$ | $(5.62\times10^{-9})$ | $(7.60\times10^{-9})$ |
| $\sigma^2$ | $2.61\times10^{-7}$ | $8.41\times10^{-8}$ | $1.51\times10^{-7}$ |
| | $(1.00\times10^{-9})$ | $(4.19\times10^{-10})$ | $(6.15\times10^{-10})$ |
| $D$ | $1.35\times10^{-3}$ | $4.84\times10^{-4}$ | $8.13\times10^{-4}$ |
| | $(7.24\times10^{-6})$ | $(3.81\times10^{-6})$ | $(4.98\times10^{-6})$ |
| *Simulation* 3 | | | |
| $\beta$ | $8.10\times10^{-7}$ | $3.02\times10^{-7}$ | $4.94\times10^{-7}$ |
| | $(5.60\times10^{-9})$ | $(2.55\times10^{-9})$ | $(3.60\times10^{-9})$ |
| $\sigma^2$ | $3.53\times10^{-8}$ | $1.00\times10^{-8}$ | $1.96\times10^{-8}$ |
| | $(1.45\times10^{-10})$ | $(3.60\times10^{-11})$ | $(7.30\times10^{-11})$ |
| $D$ | $2.76\times10^{-4}$ | $9.67\times10^{-5}$ | $1.64\times10^{-4}$ |
| | $(2.06\times10^{-6})$ | $(9.58\times10^{-7})$ | $(1.35\times10^{-6})$ |

Table S2: Mean absolute difference in the estimates produced by lmer and BLMM for $\beta, \sigma^2$ and $D$, averaged across voxels and simulation instances. Each reported average is an image-wide mean, taken across approximately $218,000$ voxels, further averaged across 1000 simulation instances. In each simulation instance, the model employed included 500 observations generated according to the methods outlined in Section 2.2. Empirical standard errors, taken across simulation instances, are also given in brackets underneath each entry in the table.

#### S9 Mean Absolute Difference for Parameter Estimation (n=1000)

| Method | Missing-data voxels | Full-data voxels | All voxels |
|---|---|---|---|
| *Simulation* 1 | | | |
| $\beta$ | $6.33 \times 10^{-10}$ | $4.19 \times 10^{-10}$ | $5.05 \times 10^{-10}$ |
| | $(7.66 \times 10^{-13})$ | $(5.85 \times 10^{-13})$ | $(6.20 \times 10^{-13})$ |
| $\sigma^2$ | $1.57 \times 10^{-10}$ | $1.10 \times 10^{-10}$ | $1.29 \times 10^{-10}$ |
| | $(8.10 \times 10^{-14})$ | $(5.84 \times 10^{-14})$ | $(5.15 \times 10^{-14})$ |
| $D$ | $4.96 \times 10^{-7}$ | $2.49 \times 10^{-7}$ | $3.48 \times 10^{-7}$ |
| | $(5.66 \times 10^{-10})$ | $(3.38 \times 10^{-10})$ | $(3.91 \times 10^{-10})$ |
| *Simulation* 2 | | | |
| $\beta$ | $6.54 \times 10^{-8}$ | $1.20 \times 10^{-8}$ | $3.36 \times 10^{-8}$ |
| | $(2.95 \times 10^{-10})$ | $(4.35 \times 10^{-11})$ | $(1.33 \times 10^{-10})$ |
| $\sigma^2$ | $6.71 \times 10^{-9}$ | $1.24 \times 10^{-9}$ | $3.46 \times 10^{-9}$ |
| | $(1.55 \times 10^{-11})$ | $(1.45 \times 10^{-12})$ | $(5.93 \times 10^{-12})$ |
| $D$ | $6.25 \times 10^{-5}$ | $1.65 \times 10^{-5}$ | $3.51 \times 10^{-5}$ |
| | $(3.32 \times 10^{-7})$ | $(9.35 \times 10^{-8})$ | $(1.80 \times 10^{-7})$ |
| *Simulation* 3 | | | |
| $\beta$ | $5.96 \times 10^{-8}$ | $2.66 \times 10^{-8}$ | $3.99 \times 10^{-8}$ |
| | $(3.10 \times 10^{-10})$ | $(1.32 \times 10^{-10})$ | $(1.71 \times 10^{-10})$ |
| $\sigma^2$ | $5.80 \times 10^{-9}$ | $3.78 \times 10^{-9}$ | $4.60 \times 10^{-9}$ |
| | $(1.29 \times 10^{-11})$ | $(6.80 \times 10^{-12})$ | $(7.17 \times 10^{-12})$ |
| $D$ | $2.97 \times 10^{-5}$ | $2.07 \times 10^{-5}$ | $2.44 \times 10^{-5}$ |
| | $(1.02 \times 10^{-7})$ | $(3.29 \times 10^{-8})$ | $(5.74 \times 10^{-8})$ |

Table S3: Mean absolute difference in the estimates produced by lmer and BLMM for $\beta, \sigma^2$ and $D$, averaged across voxels and simulation instances. Each reported average is an image-wide mean, taken across approximately $218,000$ voxels, further averaged across 1000 simulation instances. In each simulation instance, the model employed included 1000 observations generated according to the methods outlined in Section 2.2. Empirical standard errors, taken across simulation instances, are also given in brackets underneath each entry in the table.

#### S10 Mean Absolute Difference for Maximized ReML Criteria

| Method | Missing-data voxels | Full-data voxels | All voxels |
|---|---|---|---|
| *Simulation* 1 | | | |
| $n = 200$ | $8.72 \times 10^{-10}$ | $6.16 \times 10^{-10}$ | $7.02 \times 10^{-10}$ |
| | $(2.98 \times 10^{-12})$ | $(2.23 \times 10^{-12})$ | $(2.21 \times 10^{-12})$ |
| $n = 500$ | $6.24 \times 10^{-10}$ | $7.36 \times 10^{-10}$ | $6.95 \times 10^{-10}$ |
| | $(1.87 \times 10^{-11})$ | $(1.82 \times 10^{-11})$ | $(1.32 \times 10^{-11})$ |
| $n = 1000$ | $1.11 \times 10^{-9}$ | $8.23 \times 10^{-10}$ | $9.36 \times 10^{-10}$ |
| | $(4.92 \times 10^{-11})$ | $(6.02 \times 10^{-11})$ | $(4.04 \times 10^{-11})$ |
| *Simulation* 2 | | | |
| $n = 200$ | $4.25 \times 10^{-3}$ | $2.42 \times 10^{-3}$ | $3.05 \times 10^{-3}$ |
| | $(4.14 \times 10^{-5})$ | $(3.04 \times 10^{-5})$ | $(3.39 \times 10^{-5})$ |
| $n = 500$ | $2.03 \times 10^{-4}$ | $6.96 \times 10^{-5}$ | $1.20 \times 10^{-4}$ |
| | $(2.43 \times 10^{-6})$ | $(1.28 \times 10^{-6})$ | $(1.63 \times 10^{-6})$ |
| $n = 1000$ | $1.02 \times 10^{-5}$ | $2.46 \times 10^{-6}$ | $5.58 \times 10^{-6}$ |
| | $(3.99 \times 10^{-7})$ | $(2.34 \times 10^{-7})$ | $(2.12 \times 10^{-7})$ |
| *Simulation* 3 | | | |
| $n = 200$ | $1.11 \times 10^{-3}$ | $6.72 \times 10^{-4}$ | $8.23 \times 10^{-4}$ |
| | $(1.29 \times 10^{-5})$ | $(9.51 \times 10^{-6})$ | $(1.05 \times 10^{-5})$ |
| $n = 500$ | $2.55 \times 10^{-4}$ | $1.75 \times 10^{-4}$ | $2.05 \times 10^{-4}$ |
| | $(2.61 \times 10^{-6})$ | $(1.87 \times 10^{-6})$ | $(1.74 \times 10^{-6})$ |
| $n = 1000$ | $5.78 \times 10^{-5}$ | $3.43 \times 10^{-5}$ | $4.38 \times 10^{-5}$ |
| | $(1.22 \times 10^{-6})$ | $(9.44 \times 10^{-7})$ | $(7.58 \times 10^{-7})$ |

Table S4: Mean absolute difference in the maximized ReML criteria produced by lmer and BLMM, averaged across voxels and simulation instances. Each reported average is an image-wide mean, taken across approximately $218,000$ voxels, further averaged across 1000 simulation instances. The number of observations present in the model used for each simulation setting is displayed on the left. All observations were generated according to the methods outlined in Section 2.2. Empirical standard errors, taken across simulation instances, are also given in brackets underneath each entry in the table.
